# Discovering subclinical effects of limited outdoor access on gait and hoof health of cows housed in movement-restricted environments

**DOI:** 10.64898/2026.01.28.702303

**Authors:** Shabnaz Mokhtarnazif, Amir Nejati, Elise Shepley, Gabriel M. Dallago, Abdoulaye B. Diallo, Elsa Vasseur

## Abstract

Most common housing systems for dairy cows restrict their movement, which can influence welfare, gait, and hoof health of dairy cows. Outdoor access has been proposed as a management practice to offset these restrictions, but reported effects on cows’ locomotion vary and may not always be captured by traditional clinical assessments. In this study, we investigated gait and hoof through clinical (i.e., visual locomotion scoring and hoof lesion assessment) and subclinical (3D motion analysis, kinetic assessment, hoof infrared thermography and measuring claw conformation) methods to assess how limited provision of outdoor access affects non-lame cows housed in movement restricted environment. Thirty-six Holstein tie-stall cows were either given 1day/week (EX1) or 3days/week (EX3) of outdoor access (1h/day) during 5 consecutive weeks. Clinical and subclinical assessments of gait and hoof were performed before (Pre-trial), after 5 weeks of outing (Post-trial) and 8 weeks after outing (Follow-up). The results of this study revealed no clinical effect of outdoor access on cows’ locomotion score and hoof lesion prevalence. However, for subclinical assessment, both groups showed an increase in stride and stance time at Post-trial, with an increase in pressure applied by cows while standing in EX3 group and a reduction in coronary band temperature in both groups at Post-trial and Follow-up. Contact area and claw conformation changed after provision of outdoor access in both groups. This study illustrates that with the use of subclinical methods; we can reveal effects of outdoor access on gait and hoof health that might not be visible using the traditional methods.

## 1. Introduction

The increase in global population has led the dairy industry to move cows from pastures to indoor housing systems as a means of intensification. Limited land use, the ability to consistently provide high-quality feed for cows, and the control over their environment in indoor housing systems offer farmers the opportunity to expand their farms and satisfy the increasing demand for dairy products (Arnott et al., 2017). The increase intensification and indoor housing systems can compromise the welfare of dairy cows including limitation in their movement ability and increasing the risk of health issues such as lameness and mastitis (Barkema et al., 2015, Arnott et al., 2017). In Canada, dairy farms predominantly use indoor housing systems, with 39.1% employing free stalls and 59.4% using tie-stalls (C.D.I.C., 2023). Tie-stalls are the most restrictive housing systems for dairy cows in which they are confined and do not have the ability to fulfill their natural behavioral needs (Shepley et al., 2020).

One solution to mitigate the adverse effects of indoor housing is the partial provision of outdoor access. Studies showed that access to outdoor can increase locomotor activity in cows (Shepley et al., 2020, Cellier et al., 2025) and lower risk of lameness and leg injuries (Palacio et al., 2023, Blaga Petrean et al., 2024). Provision of outdoor has been associated with a lower rate of health issues such as mastitis and risk of culling (Blaga Petrean et al., 2024), better human-animal relationships and more pronounced positive emotions such as being more active, content and relaxed compared to cows with no access to outdoor (Aigueperse and Vasseur, 2021, Blaga Petrean et al., 2024). Due to the proven positive effects of outdoor access on the welfare of cows, provision of outdoor access for exercise and freedom of movement is now mandatory under the Canadian Code of Practice for the Care and Handling of Dairy Cattle during the production cycle of tethered Canadian dairy cows (NFACC, 2023).

Housing type, the opportunity of movement and cows’ fitness can directly affect cows’ gait and their susceptibility to lameness, which is defined as any deviation from normal gait or posture due to a painful condition (Van Nuffel et al., 2015). Lameness is a major welfare issue and one of the leading causes of culling, with significant economic losses in dairy farms (Huxley, 2013, Vogel et al., 2018). Locomotion scoring systems have been developed to assess lameness severity in dairy cows based on specific gait traits, where the sequence and extent of changes in these traits are associated with variations in overall gait scores (Sprecher et al., 1997, Flower and Weary, 2006, Shepley and Vasseur, 2021a). Clinical lameness is characterized by overt signs such as limping, an arched back, and reluctance to bear weight on the affected limb. This level of lameness is often accompanied by visible hoof lesions or higher locomotion scores when using traditional visual locomotion scoring systems. Some of these systems using 5-point scale (Sprecher et al., 1997, Flower and Weary, 2006) where 1 indicates normal gait and 5 indicates severe lameness. Traditional visual locomotion scoring systems, however, are limited in their ability to detect mild or subtle gait impairments (Schlageter-Tello et al., 2014, Van Nuffel et al., 2015), which may be classified as subclinical lameness. According to Merriam-Webster (2024), subclinical is defined as “not detectable or producing effects that are not detectable by the usual clinical tests”. In this context, cows at early stages of lameness may only exhibit minor deviations in gait (e.g., a locomotion score 3 and lower) and lack visible hoof lesions.

Gait and lameness assessment in cows can be performed at two levels: clinical, where visible signs are detected through human observation methods such as visual locomotion scoring and examination of claw lesions during hoof trimming (Flower et al., 2005); and subclinical, which involves the use of advanced technologies and specialized tests to identify subtle signs that are not immediately visible (Rosamond and Couper, 2022). Various technologies have been used to assess gait, which can be categorized in three main groups: kinematic gait assessment, kinetic gait assessment, and accelerometry (Bradtmueller et al., 2023). Technology used in kinematic gait assessment focuses on joint motions, limb segment angles, segment orientations and cows’ posture while walking with the aim of enabling automatic lameness detection (Alsaaod et al., 2019, Zhong et al., 2021). Vision-based technologies are the primary tools used in kinematic gait assessment; however, data from force and pressure platforms, as well as accelerometers, have also been employed to evaluate specific gait variables such as walking speed, stride length, and stance time (Bradtmueller et al., 2023, Nejati et al., 2023). Kinetics, defined as a subdivision of biomechanics, studies the forces and torques applied during a walk or a standing period (Hall, 2019). Force and pressure platforms alongside weight distribution platforms have been used broadly to assess the kinetic variables in cows’ gait (Bradtmueller et al., 2023, Nejati et al., 2023). Accelerometry approaches to investigate the acceleration data of the cow’s movement in 1-3 dimensions, usually using an accelerometer attached to the cow’s body (e.g., leg or neck) to look for the changes or abnormalities in movement behavior (Alsaaod et al., 2019, Nejati et al., 2023).

Alongside gait-related technologies, thermography is another tool used to investigate the presence of heat associated with pain or inflammation before clinical signs appear (Alsaaod et al., 2015). Infrared thermography (IRT) is known as an accurate and non-invasive technique that converts the electromagnetic radiation of objects warmer than absolute zero into false color or gray scale images (Speakman and Ward, 1998, Casas-Alvarado et al., 2020). They can detect changes in the temperature two weeks before the clinical signs appear (Schaefer et al., 2004). Studies showed that changes in cows’ hoof temperature have been associated with higher locomotion scores and incidence of claw lesions such as digital dermatitis (Alsaaod et al., 2015, Rodríguez et al., 2016).

Although some studies showed a positive effect on lameness reduction after the provision of outdoor access (Popescu et al., 2013, Palacio et al., 2023), other studies found no significant effects on cows’ gait (Loberg et al., 2004, Chapinal et al., 2010). These inconsistencies may stem from variations in housing types, frequency and duration of outdoor access (Shepley et al., 2020) and severity of lameness. Additionally, the effect might be too subtle to be detected through clinical assessments. For example, Nejati et al. (2024) showed 1-point reduction in the overall gait score of tie-stalled cows that had access to the outdoor for 5 days per week over 5 consecutive weeks compared to those without such access, although the difference was not statistically significant.

The objective of this study was to integrate technology (i.e., sub-clinical) with human (clinical) assessments to investigate the effects of low-frequency outdoor access on the gait and hoof health of non-clinically lame cows housed in movement-restricted systems. We hypothesized that this limited outdoor access would yield only subclinical effects on gait and hoof health, which could be detected using the technologies.

## 2. Materials and Methods

This experiment was conducted from October 2021 to February 2022 at Dairy Complex of McGill University and all procedures and animal use were approved by the Animal Care Committee of McGill University and affiliated hospitals and research institutes (Protocol #2016-7794).

Thirty-six lactating Holstein cows housed in a tie-stall barn were selected for this study. They were placed in 4 adjacent rows and milked every 12 hours. Cows were grouped into 6 blocks

based on the days in milk (DIM) and parity. Cows of each block were randomly assigned to one of the two treatments: 1 day/week (EX1) or 3 days/week (EX3) of outdoor access (i.e., 3 cows/treatment/block). The trial was conducted over 5 consecutive weeks. On the outing days, from 1000 to 1100, cows had access to exercise yard covered with grass and snow due to the weather condition with no potential for grass regrowth. The exercise yard is located adjacent to the barn and was equally divided into 6 paddocks of 117 m^2^. On the first week of outing, each block was randomly assigned to a paddock and rotated clockwise each week. Clinical and subclinical assessment of gait and hoof health were done at 3 data collection points (Figure 1): Pre-trial (i.e., before the trial), Post-trial (i.e., at the end of the trial) and Follow-up (i.e., 8 weeks after the trial).

**Figure 1.**
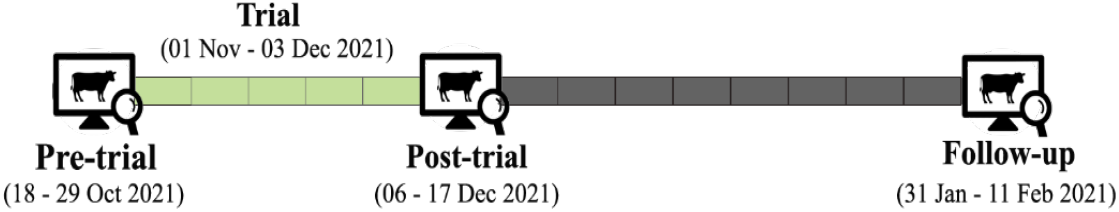
This figure illustrates the timeline of the experiment and 3 data collection points

### 2.1. Gait Analysis

Cows’ gait analysis was conducted at the three data collection points using both clinical and subclinical assessments. Because of logistic constraints, a subsample of 22 cows (i.e., 11 cows from EX1 and 11 from EX3) were selected for gait analysis at the three data collection points based on their previous exposure to the walking setting and ease of handling; however, during the Follow-up, one of the cows was culled and 21 cows were assessed.

#### 2.1.1. Clinical Assessment: Visual Locomotion Scoring

For visual locomotion scoring, a trained handler walked cows in a 5-meter test corridor with concrete flooring covered with rubber mats to record a successful gait passage (i.e., cow walked in straight line with a consistent pace and no running, stopping, defecating, or urinating). At each data collection point, cows walked for a maximum of 5 minutes. If a usable passage was not obtained, the cow was returned to her stall for the day and tested again on the last day of the collection point. The passage was considered missing data if it did not meet the listed criteria. To encourage cows to walk, when needed, the handler or another person carried a bucket filled with grain 1 meter in front of the cow while she was walking. Additional handlers were used occasionally to facilitate straight-line walking and maintain a consistent speed.

The same recorded passages were used for both visual gait scoring and subsequent 3D motion analysis. Cows’ walking was recorded using high-performance cameras (Basler Ace, Ahrensburg, Germany) fixed to the ceiling with a perpendicular view of the corridor (one from the right side; one from the rear). A total of 65 passages (22 recordings from pre-trial, 22 from post-trial, and 21 from follow-up) were recorded, resulting in 130 recordings (65 passages x 2 views recorded/passage). To blind the trained scorer to the data collection point and cows’ treatment, all the recordings were coded with numbers and randomized. Shepley and Vasseur (2021) method was used to score gait attributes, including back arch, tracking-up, joint flexion, asymmetric gait, and reluctance to bear weight using the side-view camera and swinging out using the rear-view camera due to the better vision, from 0-5 with 0.5 intervals (Table 1). The overall gait score was assigned to cows on a numerical rating system (NRS) based on Flower and Weary (2006) method and their attribute scores (Nejati et al., 2024). Cows were categorized as non-lame (NRS < 3), moderately lame (NRS = 3), lame (NRS = 4), and severely lame (NRS = 5). The weighted kappa inter- and intra-observer reliability was calculated 0.82 and 0.64, respectively, for the overall gait score and 0.39 and 0.37 for the six gait attributes.

**Table 1.** Gait attribute definitions and their associated ratings (0-5, described by Shepley and Vasseur (2021a) and adapted from (Flower and Weary, 2006))

| Gait Attribute | Definition | Scoring Scale with 0.5 intervals |  |
| --- | --- | --- | --- |
|  |  | 0 | 5 |
| Swinging out | Description of visual gait variables and the corresponding endpoints of a visual analog scale | Hind legs moving in straight line during the swing phase | Pronounced, circular motion of the hind legs during the swing phase |
| Arch back | The shape of the spine when the cattle walks | Flat spine | Convex arch between the withers and tailbone |
| Tracking up | The gap between the imprint left behind the front hoof and the new imprint formed from the rear hoof. | Hind hoof falls in imprint left by the front hoof of the same side | Hind hoof falls short of the imprint left by the front hoof of the same side |
| Joint flexion | Related to the flexions and extensions of the limb while the cow is moving | All limbs flex and extend easily | All limbs are stiff and limited in their range of motion |
| Asymmetric step | How even the stepping pattern of a cow is | Equal steps: cow places her hooves in an even “1, 2, 3, 4” rhythm | Not equal; cow places her hooves in an uneven rhythm |

#### 2.1.2. Subclinical Assessment: 3D motion Analysis

A comprehensive 3D motion analysis was performed for subclinical assessment of gait and obtain detailed measurements of different gait attributes. Prior to walking cows in the test corridor, 20 reflective markers were attached to the anatomical landmarks of cows. Sixteen markers attached to cow’s legs (i.e., 4X Coffin joints, 4X fetlocks, 2X carpals, 2X hocks, 2X elbows, and 2X stifles) and 4 on her back (i.e., 1X withers, 1X on the Spinous process of the 13^th^spinal cord, 1X on the Sacro-lumbar joint and 1X on the Sacrococcygeal joint). Reflective markers were attached to cows by first placing duct tape on the designated anatomical landmarks, followed by securing the markers on them using double-sided tape. The tail switch was restrained with a cloth strap to prevent marker obstruction during walking.

For 3D motion analysis, cows’ passages were recorded using 6 high-performance cameras (Basler Ace, Ahrensburg, Germany) attached to the ceiling (i.e., 3 cameras on the right and 3 cameras on the left side of the test corridor) at 2.3 m from the ground such that each camera view overlapped with at least 2 other camera views. All cameras were wired to a desktop computer, and videos were captured using CONTEMPLAS TEMPLO capture engine (CONTEMPLAS GmbH, Kempten, Germany) to ensure synchronization of all videos. Cameras were set to 60 fps, with 350 shutters per second and a gain of 75. The image size and position of the area of interest were 1280 width x 800 height pixels. To ensure the accuracy of data collected from 3D motion analysis, before the first recording of each test day, a calibration device with 24 spherical reflective markers was placed in the middle of the test corridor (volume dimension: 196,84 cm x 196,84 cm x 196,84 cm) and positioned such that all 24 markers were visible by all 6 cameras to record the calibration videos.

Videos were analyzed using the Vicon Motus (Version 10.0.1, Vicon motion systems Inc. Oxford, UK) software. In the first step, marker positions, definitions of angles based on the marker positions (e.g., hock angle defined as the angle made by fetlock, hock, and stifle markers), and events (i.e., the time of start and end points of each passage, toe-off and landing of each limb) were labelled in the software. In the next step, the markers from the calibration video for each day were tracked, and then 6 videos of each passage were exported to the software. A passage was defined as the time a cow started to walk in a straight line in the test corridor to the time she left the corridor, which consisted of 2 to 3 full strides. Based on the targeted traits, only 11 markers out of 20 needed to be digitized and tracked using Motus Software for each passage. These included markers attached to the coffin joints of all four legs (4 markers), fetlock, hock, and stifle joints of the rear legs (6 markers), and the one on the withers (1 marker). This enabled the calculation of track-up (cm), stride length (cm), stride time (s), velocity (cm/s), and the hock joint’s range of motion for each passage (Table 2). The coordinates of each marker represented the X (horizontal longitudinal axis corresponding to the cow’s walking direction), Y (horizontal transverse axis) and Z (vertical) axis. The data from withers’ marker was used to determine cow’s walking direction and to align the X-axis accordingly. For this purpose, the calibration frames were rotated 15-25 degrees around the Z-axis prior to data extraction. The hock angles were calculated using the Motus software, and the other variables were calculated using the coordinates of coffin markers at each event.

**Table 2.** The kinematic variables, their definitions and the method used to calculate them based on the data provided by the Vicon Motus software (Version 10.0.1, Vicon motion systems Inc. Oxford, UK)

| Variable<sup>1</sup> | Unit | Definition |
| --- | --- | --- |
| Stride length | cm | The average distance between the two consecutive stride strikes (coordinate of coffin marker at the toe-on event) of the same limb in XY axis |
| Stride time | s | The number of frames between two consecutive toe-on events of the same hoof ÷ 60 |
| Velocity | cm/s | Stride length ÷ stride time |
| Stance time | s | The number of frames between toe-on (when the hoof meets the ground) and the next toe-off (the moment the hoof leaves the ground) ÷ 60 |
| Track upX | cm | The average distance between the coordinates of coffin marker of fore limb at toe-on events and coffin marker of the ipsilateral rear limb in X axis (longitudinal axis, or the cow's walking direction) |
| Track upXY | cm | The average distance between the coordinates of coffins markers of fore limb at toe-on event and coffin marker of the ipsilateral rear limb in XY axis (the diagonal line connecting longitudinal (X) and Transverse (Y) axis) |
| Hock angle (Min) | degree | The lowest number recorded from the angle created between the fetlock, hock, and stifle markers during walking from the first toe-on to the last toe-off |
| Hock angle (Max) | degree | The highest number recorded from the angle created between the fetlock, hock, and stifle markers during walking from the first toe-on to the last toe-off |
| Hock angle range of motion (ROM) | degree | The difference between Maximum and Minimum hock angles |
| Hock angle (Med) | degree | The median of the numbers recorded from the angle created between the fetlock, hock, and stifle markers during walking from the first toe-on to the last toe-off of each passage |
| Hock angle (Avr) | degree | The average of hock angle at all toe-on and toe-off events of each passage |

<sup>1</sup>Abbreviations: Min = Minimum; Max = Maximum; Med = Median; Avr = Average

#### 2.1.3. Subclinical Assessment: Kinetic Assessment

Kinetic analysis of gait, which serves as another tool for subclinical and objective measurement of gait, was conducted at all data collection points (i.e., Pre-trial, Post-trial, and Follow-up) on the subsample of cows. At each data collection point, cows were individually tied in the kinetic pen such that she was standing on 2 pressure plates (FootScan®, Teckscan Inc. MA, USA) parallel to one another with their left legs (front and hind) on one plate and the right legs (front and hind) on the other. Plates were connected to two portable laptops, and the FOOTSCAN software (FootScan®, Teckscan Inc. MA, USA) was used to visualize and capture the data while the cow stood still on the plates. In the first step, a screenshot (SC) per plate of the images visualized by the FOOTSCAN software was taken through the software, then data from pressure plate was recorded with 125 frame per seconds for 30 seconds (30Srec) for both plates at the same time. This procedure was successful if cows had their hoof prints fully on the plates without any movement during the 30Srec otherwise it was repeated.

Different kinetic variables were measured depending on the data collection strategy (SC vs 30Srec). Contact area (cm^2^), as well as maximum and average pressure (N/cm^2^) were measured using both SC and 30Srec. Maximum and average force (N) were only measured using 30Srec, whereas pressure distribution ratio was measured using SC data only. Using 30Srec, force and pressure were calculated by the software at each frame based on the footprint zones. Since the software is designed based on human anatomy, we first defined each claw as separate zones and then calculated the maximum and average force (N) as well as pressure (N/cm^2^) measured by the software from all frames in that specific zone. To measure the contact area, pictures of SC and 30Srec were used and exported to the Adobe Photoshop software (Photoshop 2022, version 23.2, Adobe Systems Inc., San Jose, USA). The number of pixels representing the known distance in cm was calculated using the scale provided by the FOOTSCAN software (FootScan®, Teckscan Inc. MA, USA), and the contact area was calculated using the number of pixels in the hoof print area (Table 3).

**Table 3.**
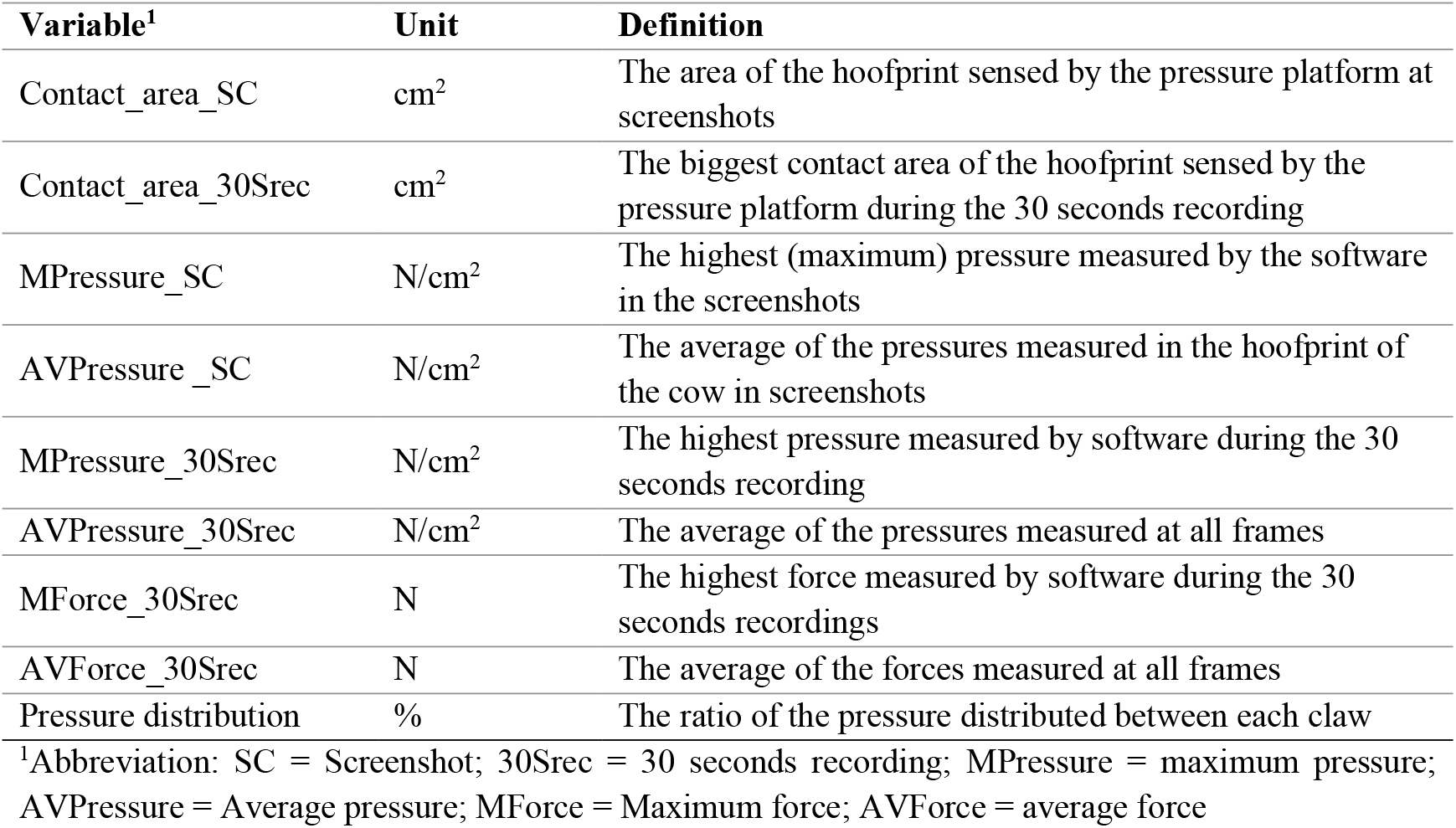
The kinetics variables and how they were calculated using the data from Footscan software (FootScan®, Teckscan Inc. MA, USA)

| **Variable<sup>1</sup>** | **Unit** | **Definition** |
| --- | --- | --- |
| Contact_area_SC | cm<sup>2</sup> | The area of the hoofprint sensed by the pressure platform at screenshots |
| Contact_area_30Srec | cm<sup>2</sup> | The biggest contact area of the hoofprint sensed by the pressure platform during the 30 seconds recording |
| MPressure_SC | N/cm<sup>2</sup> | The highest (maximum) pressure measured by the software in the screenshots |
| AVPressure_SC | N/cm<sup>2</sup> | The average of the pressures measured in the hoofprint of the cow in screenshots |
| MPressure_30Srec | N/cm<sup>2</sup> | The highest pressure measured by software during the 30 seconds recording |
| AVPressure_30Srec | N/cm<sup>2</sup> | The average of the pressures measured at all frames |
| MForce_30Srec | N | The highest force measured by software during the 30 seconds recordings |
| AVForce_30Srec | N | The average of the forces measured at all frames |
| Pressure distribution | % | The ratio of the pressure distributed between each claw |

<sup>1</sup>Abbreviation: SC = Screenshot; 30Srec = 30 seconds recording; MPressure = maximum pressure; AVPressure = Average pressure; MForce = Maximum force; AVForce = average force

### 2.2. Hoof Health Assessment

Cows were restrained in an upright hoof trimming chute for hoof health assessment, including clinical hoof health assessment, sole thermography and measuring claw conformation. A full hoof trimming using the five-step Dutch method (Toussaint, 1989) was conducted a month before the start of the Pre-trial point. In order to clean hooves before our measurements, sliver horn trimming - in which a thin or “sliver” layer of the sole is removed to clean the sole and evaluate hoof health and conformation without significantly altering the hoof’s overall structure - was performed at the Pre-trial, Post-trial and Follow-up points. A full hoof trimming was conducted after the sliver trimming at follow-up (8 weeks after trial) to investigate possible hoof lesions and pathologies.

#### 2.2.1. Clinical Assessment: Hoof Lesion Assessment

Hoof health was clinically assessed for all 36 cows at the three data collection points, however only data from the Pre-trial and Follow-up was used for analysis because at Post-trial, no infectious lesions were recorded and non-infectious lesions at this time were considered unreliable indicator of the outdoor access effects due to the 8-week development period required for claw horn disruption lesions to manifest the effects of metabolic and mechanical stress on the sole (Shearer et al., 2015). Claw lesions were recorded based on ICAR Claw Health Atlas (Egger-Danner et al., 2014) where zone 3 represents white line lesions and zone 4 represent sole ulcer. A system from the combination of Flower et al. (2006) and Nikkhah et al. (2005) were used to create a 5-point scaling system to score severity of lesions (0 = no hemorrhage and/or discoloration, 1= slight hemorrhage, 2= moderate hemorrhage, 3= severe hemorrhaged lesion and potential visibility of fresh blood, 4= exposed of the corium, ulcer).

#### 2.2.2. Subclinical Assessment: Infrared Thermography

Infrared thermography (IRT) was used for subclinical hoof health assessment using a handheld IRT camera (FLIR E8, Teledyne FLIR LLC, Wilsonville, Oregon, USA), which captured both thermal and digital images. The camera had a temperature range of –20°C to 250°C, with a thermal sensitivity of 0.05°C and an accuracy of ±2%. The lens had a 45° × 34° field of view, and the resolution was 320 × 240 pixels. Camera settings were adjusted with an emissivity of 0.95, object distance of 1 m, and reflected temperature of 20°C. IRT images were taken from each hoof from two separate views (i.e., dorsal -coronary band and plantar -sole). An IRT image of the cow’s eye was taken at each data collection to use as a control for individual variance in body temperature (Redaelli et al., 2019)

IRT images from the coronary band were taken from the sub-sample of 23 cows at three data collection points (Pre-trial, Post-trial and Follow-up). The digital format of images taken by the thermal camera was used to evaluate the foot hygiene (hoof, coronary band, and dew claw areas) using a 4-point hygiene scoring system ranging from 0 to 3 adapted from Schreiner and Ruegg (2003). In this scale, score 0 indicates a completely clean hoof; 1 means slight contamination with scattered manure; 2 denotes moderate contamination with manure covering most of the hoof and coronary band; and 3 represents heavy contamination, with the foot entirely covered in manure or dirt. Feet with scores 2 and 3 were dropped from further analysis. At each data collection point (Pre-trial, Post-trial and Follow-up), IRT images were taken immediately after kinetic recordings. To minimize thermogram artifacts, cows remained still for at least 1 minute before imaging.

Sole view thermal images were taken for all 36 cows using the same camera (FLIR E8, Teledyne FLIR LLC, Wilsonville, Oregon, USA) 2-5 minutes after sliver trimming and removal of dirt at the three data collections to remove the effect of trimming on the sole temperature. Both kinetic pen and the trimming chute were located inside the barn where they are protected from direct sunlight and wind flow. A thermometer (Mini Thermo-Anemometer, EXTECH instruments, Nashua, NH, USA) was placed close to the animal to record the ambient temperature after taking IRT pictures from each cow.

Data from the dorsal view was obtained using Therma-CAM Researcher Professional 2.10 software (FLIR Systems, Inc., Wilsonville, Oregon, USA). Four regions of interest (ROI) were defined: hoof coronary band area (CB), coronary band of lateral and medial claws (LCB and MCB, respectively; Figure 2). Maximum, minimum, average, maximum minus minimum, and standard deviation of the selected regions were extracted from IRT images.

**Figure 2.**
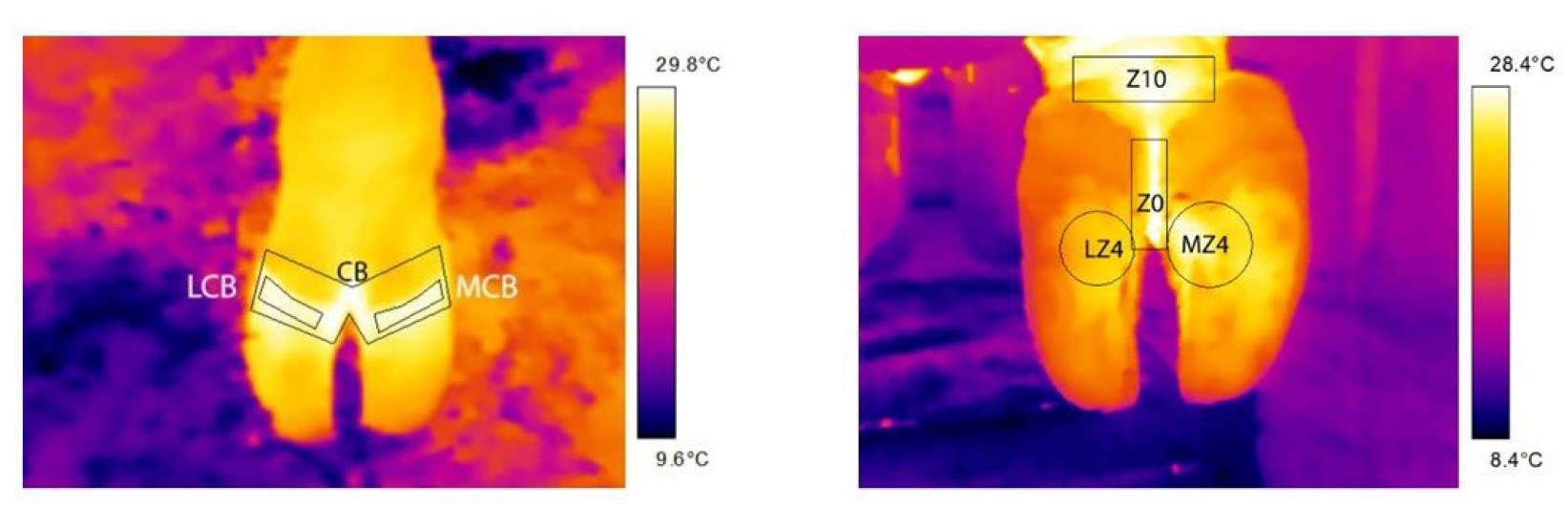
The IOR selected in dorsal and plantar views. Dorsal view: CB = hoof coronary band, MCB = medial claw coronary band and LCB = lateral claw coronary band. Plantar view: Z10 = one 10 or the skin above the claws, Z0 = Zone 0 or the skin below the claws, LZ4 and MZ4 = Zone 4 of lateral and medial claws, respectively.

To obtain sole view data, pictures were imported to the FLIR TOOLS software (Version 6.4.18039.1003, FLIR Systems, Inc., Wilsonville, Oregon, USA) and cow’s eye and four anatomical zones based on the description of Shearer et al. (2004) were selected as ROI: the skin above the claws or zone 10 (Z10), the skin between the claws or zone 0 (Z0), and the most common area for sole ulcer (Oikonomou et al., 2014), which is the zone 4 of both lateral (LZ4) and medial (MZ4) claws. Maximum, minimum, and average temperature of all ROIs were measured for further analysis (Figure 2). IRT image of the cow’s eye was taken at the same time of the collection of hooves images (both dorsal and sole view) and the maximum eye temperature was measured to be used as an indicator of the core body temperature.

#### 2.2.3. Subclinical Assessment: Measuring Claw Conformation

Claw conformation was measured at the three data collection points for all 36 cows enrolled in the study while they were restrained in the hoof trimming chute after the sliver hoof trimming. The measurements were done either live at the trimming chute or using digital pictures taken at the time to be analyzed later. For the live measurements, a trained observer measured claw length as the distance from the horn junction at the coronary band to the apex of the toe at the dorsal view using a ruler. The toe angle was measured as the angle of the dorsal border to the weight-bearing surface using a Digital sliding T-bevel (ANGLE-IZER®, General Tools & Instruments LLC, NJ, USA). Live measurements were taken only from the lateral claws of the hind legs due to time constraints, their greater weight-bearing role, and higher susceptibility to lesions compared to medial claws (Nuss and Paulus, 2006, Correa-Valencia et al., 2019).

Digital pictures were taken from lateral and medial claws of hind limbs using rear cameras of Samsung Galaxy 8 and iPhone 12 Pro max, to measure sole length and width following the procedure explained by Laven et al. (2015). The measurements were conducted using ImageJ software (U.S. National Institutes of Health, Bethesda, Maryland, USA) by calculating the number of pixels in the images. A ruler was placed above the hooves’ heels to serve as a scale.

### 2.3. Statistical Analysis

Linear mixed models were used to evaluate the effects of outdoor access on various variables. Block, treatment (EX1 and EX3), and time (Pre-trial, Post-trial, and Follow-up) were included as fixed effects as well as an interaction between treatment and time. Time was also used as a random slope to assess the impact of outdoor access over time for individual cows (Lohse et al., 2020). Cow was treated as a random intercept for visual locomotion scoring. For hoof lesion assessment, kinetic, and conformation measurements, claws nested within cows were used as a random intercept. Similarly, limb nested within cow was used as a random intercept for 3D motion analysis and thermography variables. Eye maximum temperature and ambient temperature were added as co-variables for thermography data. Statistical analyses were done using R Core Team (2022; Version 4.2.2, Vienna, Austria) with RStudio interface (version 2023.03.1, RStudio: Integrated Development for R, PBC, Boston, MA, USA) and the R package *nlme* (Pinheiro et al., 2020).

Residual analysis was conducted on all models to assess the assumptions of homoscedasticity, independence, and normal distribution of within-group errors, as well as to verify the normal distribution and independence of random effects. This graphical analysis followed the procedures outlined by (Pinheiro and Bates, 2000). Serial correlation structures were also examined to account for potential dependencies among observations over time. The correlation structures evaluated included general, autoregressive of order 1, and compound symmetry. The model selection process utilized the Akaike Information Criteria (AIC) to identify the best-fitting model for the data. Due to the complexity of the models, some models did not converge. For those cases, we removed the random slope of time for that specific model and then tested the models for the serial correlation structure (Models convergence is presented in Supplementary Table S1). To assess the statistical significance of fixed effects in the models, the ANOVA test was employed, with a significance level of α < 0.05. Estimated effects were evaluated using marginal means, and Bonferroni P-value adjustment was applied to account for multiple comparisons of means.

## 3. Results and Discussion

### 3.1. Clinical assessments

#### 3.1.1. Visual Locomotion Scoring

No cows with a locomotion score of more than 3 were enrolled in the study, and no cows got an overall gait score of 3.5 or more during the study. Only 3 cows in Pre-trial (2 in EX1 and 1 in EX3) and 1 cow from EX3 in Post-trial were moderately lame (NRS = 3) across the study. The overall NRS and 6 gait attributes did not differ from Pre-trial to Post-trial and Follow-up (P > 0.05), and there was no difference between the two treatment groups (P > 0.05), and no effect of interaction between time and treatment for overall gait score and gait attributes (P > 0.05). Cows started the trial with low overall locomotion score (2.16 ± 0.16 and 2.02 ± 0.16 for EX1 and EX3, respectively) and showed numerical reduction of 0.2 – 0.3 in EX1 and EX3, respectively at Post-trial, which was consistent eight weeks after the provision of outdoor access with the overall NRS of 1.91 ± 0.16 and 1.75 ± 0.16 for EX1 and EX3, respectively, at Follow-up (Figure 3). Track-up, joint flexion and reluctance to bear weight showed similar results, meaning non-significant reduction in their score from Pre-trial to Post-trial and Follow-up (Table S2 and Table S3 found in supplementary materials).

**Figure 3.**
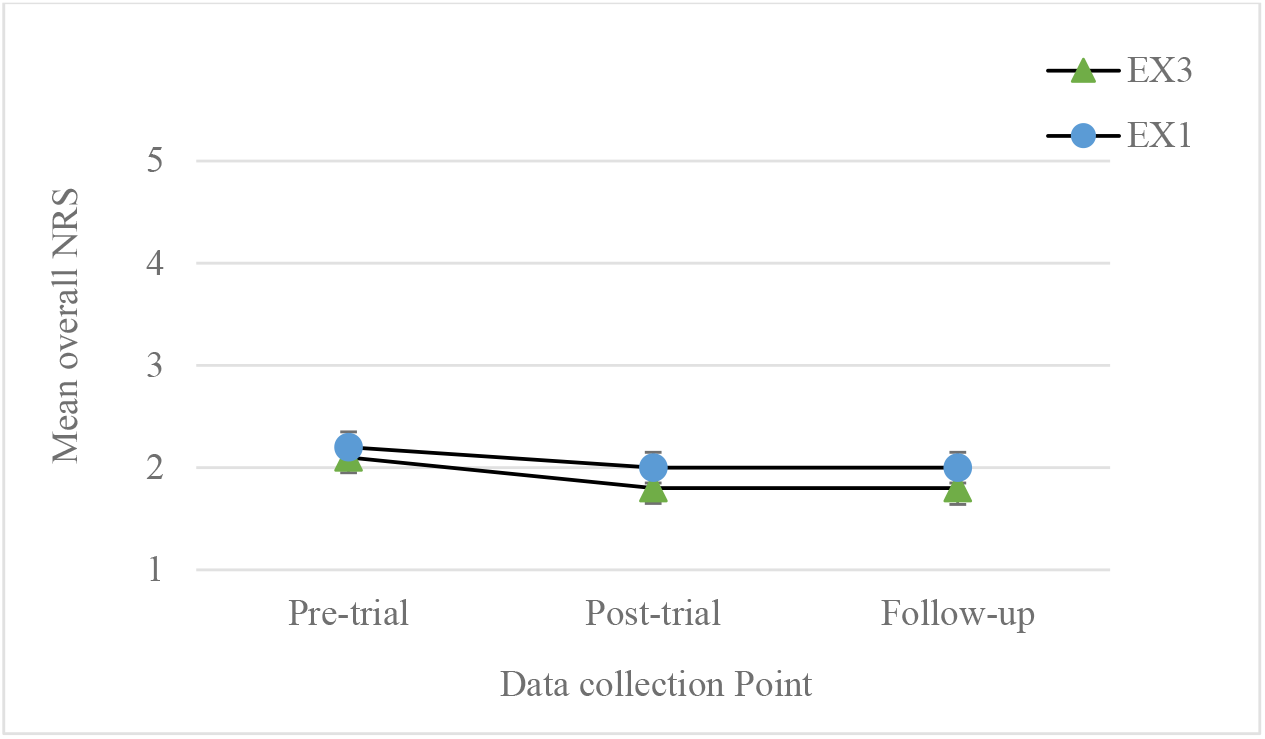
The average overall gait score (NRS) of both treatment groups at the three data collection points did not differ from Pre-trial to Post-trial and Follow-up data collection points and between treatment groups (P > 0.05).

#### 3.1.2. Hoof Lesion Assessment

A total of 41 claw lesions were recorded at the two data collection points: 22 sole hemorrhages at Pre-trial and 17 sole hemorrhages and 2 white line hemorrhages at Follow-up = 17 sole hemorrhages and 2 white line hemorrhages. Of the 39 sole hemorrhages, 30 had a severity of 1 (76.9%), 8 lesions had a severity of 2 (20.5%), and there was only one sole hemorrhage with a severity of 3 (2.6%). The two white line hemorrhages observed at Follow-up had a severity of 1 and 2 in zone 3. The prevalence of lesions was similar for EX1 (5.56 % and 6.62%) and EX3 (9.72% and 6.94%) in Pre-trial and Follow-up (P = 0.47). There was no effect of time, treatment, or their interaction on the severity of the lesions (P > 0.05).

These results showed that the limited provision of outdoor access (1h and 3h per week) would not yield any clinical effect on gait and hoof health of cows. These findings were in accordance with our hypothesis that this level of outdoor access may not be sufficient to see improvements in gait and hoof health through clinical assessment. Our results regarding the visual locomotion scoring showed no reduction in overall gait score and no improvements in any six gait attributes, which was in contrast to Nejati et al. (2024), who reported numerical reduction of 1 to 1.2 score (from a 5-point scoring system) in overall NRS and 3 gait attributes (i.e., track up, asymmetry and reluctance to bear weight) after provision of 5h/week of outdoor access in non-lame cows housed in tie-stall. Shepley and Vasseur (2021a) also showed improvement in joint flexion of non-lame tie-stalled cows after being housed in deep-bedded loose pen for 8 weeks. The initial fitness baseline and gait status have significant role in determining the effect of exercise provision. Studies indicate that outdoor access reduces lameness prevalence in herds, as observed by Corazzin et al. (2010), who found that summer grazing lowered lameness in tie-stall-housed cows. Similarly, Palacio et al. (2023) reported that limited outdoor access in winter—just a few hours weekly—resulted in less lameness compared to cows without any outdoor time. These outcomes align with earlier studies, which consistently found lower lameness levels in cows granted outdoor access across various housing systems (Bielfeldt et al., 2005, Hernandez-Mendo et al., 2007, Popescu et al., 2013).

As anticipated, claw lesion prevalence and severity remained unchanged throughout the study, paralleling findings in visual locomotion scoring. This finding aligned with those of Nejati et al. (2024), who observed no change in claw lesion prevalence or severity after five weeks of higher-frequency outdoor access (1 hour per day, 5 days a week) in cows with similar characteristics. We intentionally enrolled cows without severe lameness (only three cows at pr-trial had score 3) or ulcerative lesions to assess the impact of low-frequency outdoor access on healthy, tie-stall-housed cows unaccustomed to stall release as part of their daily routine. Increasing health issues among dairy cows are one of the main concerns raised by producers when discussing the provision of outdoor access (Smid et al., 2021). Cow-level prevalence of claw lesions has been reported at 25.7%, with a 7.1% prevalence of hemorrhage, in Canadian tie-stall dairy farms (Cramer et al., 2008). Our results showed less than 10% prevalence of sole hemorrhage. Both white line and sole hemorrhage lesions recorded in this study are considered low in severity compared to other types of claw lesions, which are usually underreported or unreported by hoof trimmers (Solano et al., 2016).

Popescu et al. (2013) found a more than 10% lower prevalence of non-infectious lesions in tie-stall cows with outdoor access compared to those without any access. As reviewed by Shepley and Vasseur (2021b), providing outdoor access is generally associated with a reduced risk of non-infectious hoof lesions. The difference with our findings may be due to the low prevalence of lesions among cows at the start of our study and the possibility that the effects of low-frequency outdoor access were too subtle to be clinically detectable.

### 3.2. Subclinical assessments

#### 3.2.1. 3D Motion Analysis

The 3D motion analysis of the gait showed that both stride and stance time changed significantly over time (P < 0.05). The average stride time for all cows showed an increase from 1.09 ± 0.01s in Pre-trial to 1.13 ± 0.01s in Post-trial and returned to the initial value of 1.09 ± 0.01s at Follow-up (P = 0.007). The average of stance time demonstrated the same pattern with an increase of 0.03s from Pre-trial to Post-trial, followed by a reduction in Follow-up (P = 0.003). No effect of time, treatment or their interaction was observed during the 3D motion analysis of gait (Table 4 and Table S4 found in supplementary materials).

**Table 4.** Average ± standard errors of gait variables obtained through 3D motion followed by the statistical significance of treatments (EX1 and EX3) and time points (i.e., Pre-trial, Post-trial and Follow-up). Different letters in superscript indicate the significance.

| Variable<sup>1</sup> | Treatment Group |  | Time |  |  | P-value |  |  |
| --- | --- | --- | --- | --- | --- | --- | --- | --- |
| | EX1 | EX3 | Pre-trial | Post-trial | Follow-up | TRT<sup>2</sup> | Time | TRT × Time |
| Stride length (cm) | $162 \pm 1.15$ | $160 \pm 1.07$ | $160 \pm 0.94$ | $162 \pm 0.96$ | $161 \pm 0.95$ | 0.31 | 0.31 | 0.54 |
| Stride time (s) | $1.10 \pm 0.01$ | $1.11 \pm 0.01$ | $1.09 \pm 0.01^a$ | $1.13 \pm 0.01^b$ | $1.09 \pm 0.01^a$ | 0.22 | 0.007 | 0.37 |
| Velocity (cm/s) | $149 \pm 1.52$ | $145 \pm 1.47$ | $148 \pm 1.47$ | $145 \pm 1.53$ | $148 \pm 1.63$ | 0.032 | 0.07 | 0.36 |
| Stance time (s) | 0.71 ±<br>0.01 | 0.72 ±<br>0.01 | 0.70 ±<br>0.01 <sup>a</sup> | 0.73 ±<br>0.01 <sup>b</sup> | 0.70 ±<br>0.01 <sup>a</sup> | 0.11 | 0.003 | 0.17 |
| Track-up XY<br>(cm) | 9.85 ±<br>0.59 | 9.43 ±<br>0.55 | 10.02 ±<br>0.57 | 9.50 ±<br>0.58 | 9.39 ±<br>0.62 | 0.48 | 0.13 | 0.08 |
| Hock angle<br>range of<br>motion (ROM) | 36.1 ±<br>0.61 | 34.9 ±<br>0.56 | 36.0 ±<br>0.63 | 34.9 ±<br>0.64 | 35.7 ±<br>0.64 | 0.82 | 0.79 | 0.30 |
| Hock angle<br>(Med<sup>1</sup>) | 147 ±<br>0.89 | 147 ±<br>0.83 | 147 ±<br>0.73 | 146 ±<br>0.75 | 147 ±<br>0.74 | 0.90 | 0.19 | 0.69 |

<sup>1</sup>Med = Median; <sup>2</sup>TRT = Treatment

Using the 3D motion analysis of gait showed temporary rise in both stride and stance time at Post-trial with the return to their initial values at Follow-up. The physical and mental satisfaction of cows during walking on suitable surface have been described as “locomotion comfort” which results in enhanced natural gait behavior (Telezhenko, 2007). The increase in the stance time (i.e., the supporting phase of the gait in which the hoof is in contact with the ground with no movement) can be a sign of locomotion comfort in cows (Alsaaod et al., 2017). For lame cows, the stance time is generally shorter in the affected limb compared to the sound limb. The reason can lie in the cow’s attempt to minimize the weight bearing on the painful leg and spend more time on the sound limbs (Kang et al., 2020). Cows with clinical lameness (score 4 and more in a 5-point scaling system) have longer stride time compared to cows with lower locomotion scores (Liu et al., 2011). However, some studies mentioned that, when cows walk on more comfortable surfaces such as rubber mats, they reduce their stride time and increase their walking speed (Flower et al., 2007, Franco-Gendron et al., 2016). Additionally, painful lesions such as sole ulcers and increased locomotion score have been reported to be associated with longer stride time and slower walking speed (Flower et al., 2007, Maertens et al., 2011). In the cited studies, the alteration in stride time was consistently linked with changes in walking speed. In contrast, our study showed an increase in stride time with no changes in walking speed or stride length across all data collection points. Swing phase compromises around two-thirds of stride time (Alsaaod et al., 2017), the increase in the stride time was predominantly driven by increase in the stance time. Therefore, while an extended stride time is generally seen as an indication of a deteriorating gait (Flower et al., 2005; Flower et al., 2007; Franco-Gendron et al., 2016), in our context, this increase was chiefly due to adjustments in the stance time, not due to changes in the swing time. We suggest that the increase in the stance time of both groups can stem from the increase in the confidence and/or locomotion comfort of the cows after 5 weeks of outdoor access. These results show that even the modest provision of outdoor access – frequencies as brief as 1 to 3 hours per week – had positive effects in cows housed in movement-restricted environments.

#### 3.2.2. Kinetic Assessment

Our results showed significant reduction in contact area from Post-trial (26.8 ± 0.62 cm^2^) to Follow-up (24.8 ± 0.67 cm^2^, P = 0.0015) from the screenshots (SC) capturing the contact area of claws on pressure plates. In recordings taken over 30 seconds (30Srec), the contact area showed a non-significant increase from Pre-trial (28.5 ± 0.69 cm^2^) to Post-trial (30.1 ± 0.66 cm^2^, P = 0.07) followed by a significant reduction to 27.5 ± 0.68 cm^2^ at Follow-up (P = 0.001) with no significant difference between the mean contact area at Pre-trial and Follow-up. There were no significant differences in mean distribution of pressure between treatment groups (EX1 and EX3) and across time points (Pre-trial, Post-trial, Follow-up), with no observable effects of treatment, time, or their interactions (Table 5).

**Table 5.** Average ± standard errors of variables obtained from pressure plate while cows stood on them for 30 seconds, followed by the statistical significance of treatments (EX1 and EX3) and time points (i.e., Pre-trial, Post-trial and Follow-up). Different letters in superscript indicate the significance

| Variable<sup>1</sup> | Treatment Group |  | Time |  |  | P-value |  |  |
| --- | --- | --- | --- | --- | --- | --- | --- | --- |
| | EX1 | EX3 | Pre-trial | Post-trial | Follow-up | TRT<sup>2</sup> | Time | TRT × Time |
| Pressure Distribution | $27.2 \pm 1.16$ | $28.1 \pm 1.15$ | $27.3 \pm 1.13$ | $28.8 \pm 1.11$ | $26.8 \pm 1.21$ | 0.70 | 0.07 | 0.11 |
| MPressure_SC | $256 \pm 15.8$ | $256 \pm 15.5$ | $234 \pm 16.0$ | $256 \pm 13.8$ | $271 \pm 21.0$ | 0.16 | 0.93 | 0.11 |
| AVPressure_SC | $41.5 \pm 2.08$ | $44.8 \pm 2.05$ | $38.3 \pm 2.12$ | $45.7 \pm 1.84$ | $45.4 \pm 2.61$ | 0.28 | 0.34 | 0.01 |
| MPressure_30Srec | 76.5 ± 4.31 | 78.5 ± 4.24 | 69.6 ± 4.36 | 79.2 ± 3.89 | 83.7 ± 4.45 | 0.11 | 0.99 | 0.04 |
| MForce_30Srec | 64.4 ± 3.73 | 68.0 ± 3.67 | 58.8 ± 3.69 | 67.9 ± 3.10 | 71.8 ± 4.35 | 0.32 | 0.88 | 0.12 |
| AVForce_30Srec | 2502 ± 144 | 2495 ± 141 | 2280 ± 180 | 2695 ± 131 | 2519 ± 147 | 0.16 | 0.33 | 0.08 |
| Contact_area_SC | 2054 ± 115 | 2167 ± 112 | 1873 ± 129 | 2301 ± 120 | 2157 ± 123 | 0.29 | 0.38 | 0.09 |
| Contact_area_30Srec | 25.8 ± 0.75 | 25.9 ± 0.75 | 26.0 ± 0.68<sup>ab</sup> | 26.8 ± 0.62<sup>a</sup> | 24.8 ± 0.67<sup>b</sup> | 0.78 | 0.02 | 0.80 |

<sup>1</sup>Abbreviations: SC = Screenshot; 30Srec= 30 seconds recording; MPressure = maximum pressure; AVPressure = Average pressure; MForce= Maximum force; AVForce = average force
<sup>2</sup>TRT = treatment

Regarding pressure, there was no significant effect of time or treatment on mean maximum and average pressure exerted by claws, whether in SC or 30Srec recordings (P > 0.05). Nonetheless, significant interactions between time and treatment were found. In SC, the average pressure for EX3 (AVPressure_SC) increased from Pre-trial to Post-trial and Follow-up (36.0 ± 3.02, 46.7 ± 2.62, and 51.8 ± 3.67 N/cm^2^, respectively, P = 0.014). Similarly, in 30Srec recordings, the maximum pressure for EX3 (MPressure_30Srec) increased from 62.7 ± 6.17 N/cm^2^ at Pre-trial to 81.6 ± 5.56 N/cm^2^ at Post-trial and 91.2 ± 6.21 N/cm^2^ at Follow-up (P = 0.04).

Although other pressure metrics (MPressure_SC and AVPressure_30Srec) did not show a significant time and treatment interaction, they did increase numerically from Pre-trial to Follow-up (Table 6). Finally, force analysis from the pressure plates showed no significant effects of treatment, time, or their interaction on the mean maximum and average force measured during the 30-second recordings (30Srec).

**Table 6.**
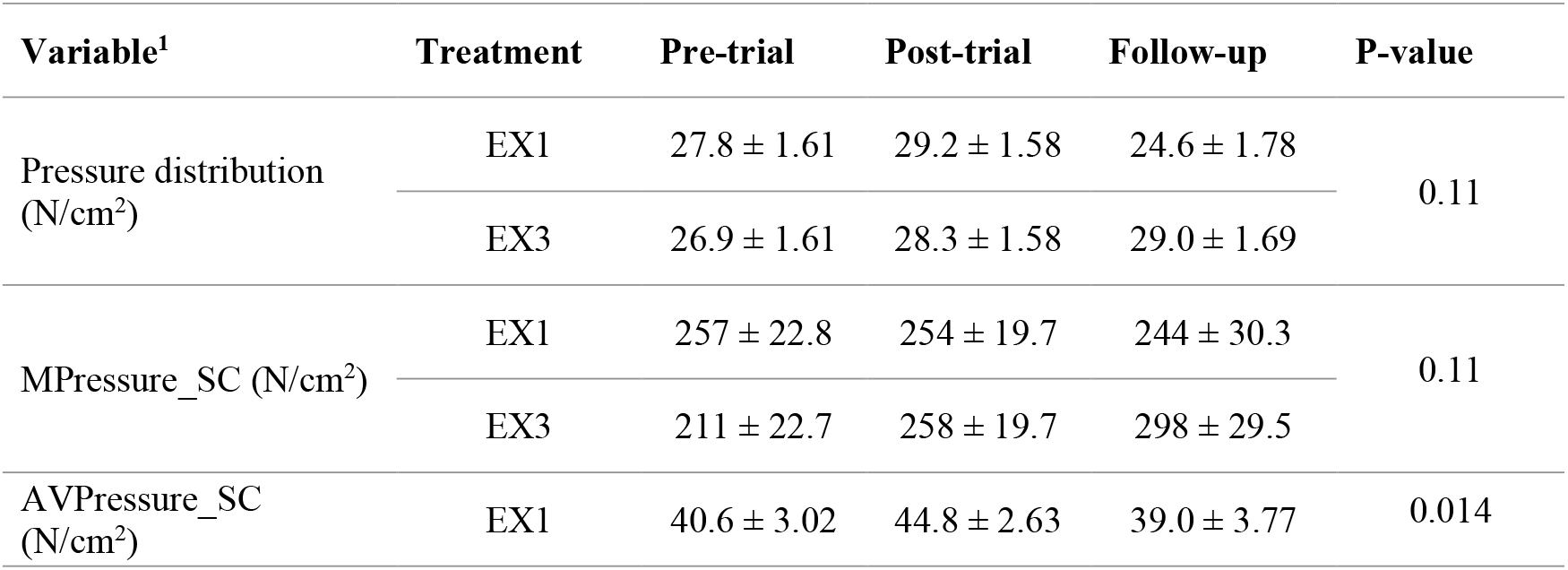
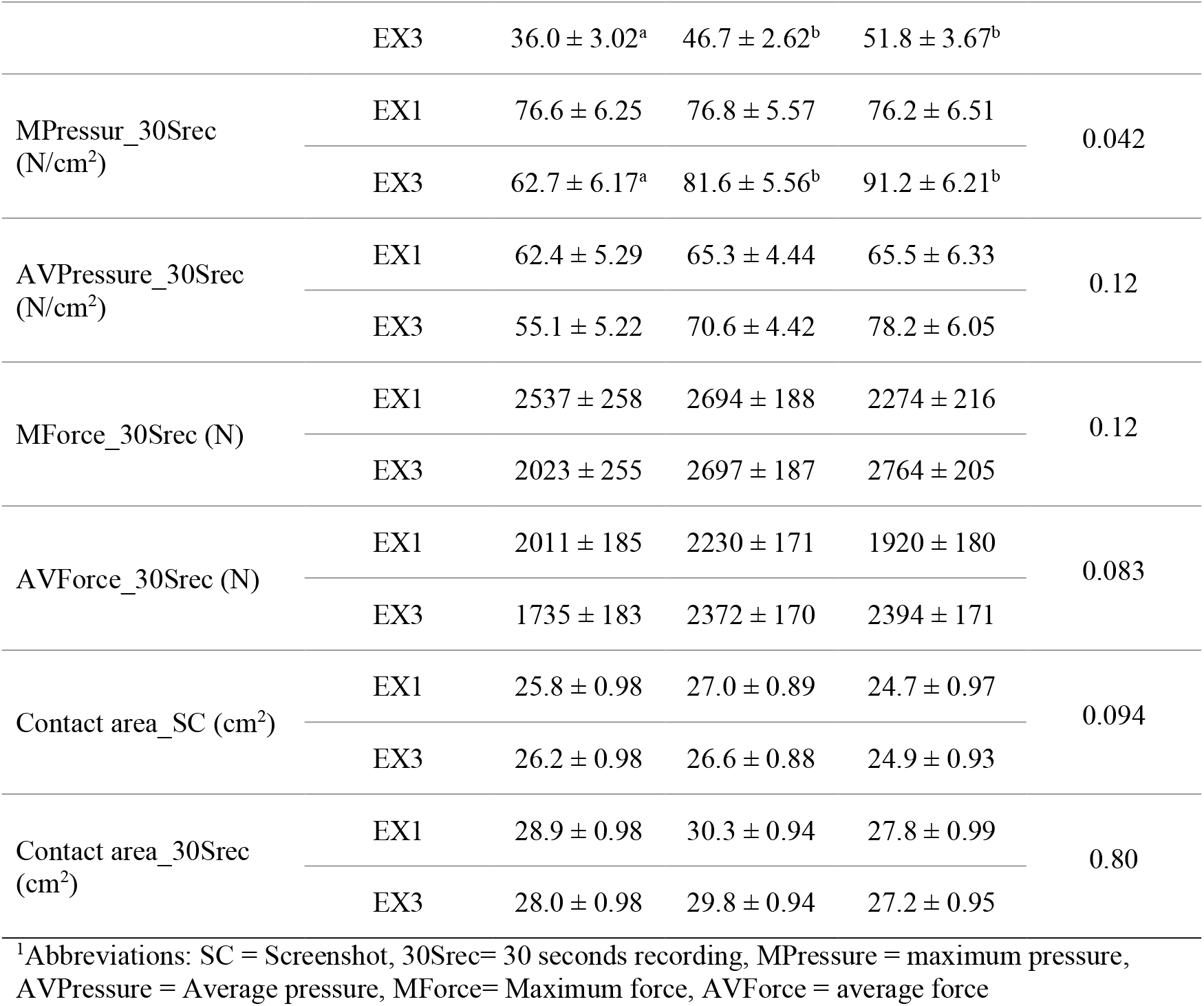
Average ± standard errors of variables obtained from the pressure plate while cows stood on them for 30 seconds based on the interaction between the two treatment groups (i.e., EX1 and EX3) and each data collection point (i.e., Pre-trial, Post-trial and Follow-up). The P-value shows the interaction effect. Different letters in superscript indicate the significance.

The kinetic assessment only performed on the rear legs and reveals an increase in the contact area at Post-trial with a drop at Follow-up, which is in accordance to the claw conformation assessment (see: Measuring Claw Conformation) in which sole length and sole width show the same pattern of increase at Post-trial, followed by a decrease at Follow-up.

Provision of outdoor access gives cows the opportunity to walk on natural surfaces (e.g., sandy alley and soil exercise yard). The increase in the contact area after 1-3h/week of outdoor access may indicate that greater movement opportunities changes the weight bearing surface due to the physical alteration in growth and wear rate of claw horn tissue (van der Tol et al., 2004, Telezhenko et al., 2008, Ouweltjes et al., 2009). When cows walk on more abrasive flooring, such as slatted concrete as opposed to rubber flooring, they display a higher contact area (Ouweltjes et al., 2009), which may contribute to a more level sole and enhanced gripping with the floor. Telezhenko et al. (2008) suggested that an increase in contact area on hard flooring might be due to disruption of horn tissue wear and expansion of contact area to the sole, which is undesirable because soles are thinner and softer and should not bear weight. However, walking on natural surfaces can develop more protruding walls to bear most of the claws’ weight (van der Tol et al., 2004). It is important to note that we did not investigate the changes in contact area in different claw zones; therefore, the area(s) where the increase has happened is unknown. After the eight weeks without outdoor access (i.e., Follow-up), we observed a reduction in contact area, possibly due to an increase in hoof growth rate resulting in longer claws and disruption in weight bearing.

Through the kinetic assessment, a notable increase in the applied pressure by the EX3 group was revealed during the Post-trial and Follow-up. We assume that there might be two reasons behind this increase. Firstly, it might be the result of observed changes in the weight-bearing surface of the claws, causing alteration in pressure distribution within the claws and leading to the concentration of pressure on specific points of the sole, which is aligned with the increase in the contact area. Secondly, the increase in the loaded pressure could be linked to the potential improvement in locomotion comfort of cows, corroborating with our results in gait and kinematic analysis. Alsaaod et al. (2017) suggested that cows tend to apply more pressure when walking on pasture during the toe-on phase in comparison to other artificial flooring. While increased pressure and force applied by claws during walking are generally considered unfavorable due to the potential risk of lesion development (Medina-González et al., 2022), it is essential to note that our study measured the applied pressure while cows were standing. Therefore, it is plausible that the observed increase in pressure is a result of cows becoming physically fitter and possessing stronger muscles after having more movement opportunities, particularly in the EX3 cows. As a complementary effect, they might be loading more pressure on their claws as they experience less pain and discomfort. For instance, Liu et al. (2011) found that cows with a locomotion score of 1 loaded more maximum and average force on the ground when compared to cows with a score of 3 or higher in a 5-point scaling system. Future research investigating changes in the contact area and pressure applied based on claw’s anatomical zones while cows are standing and walking would allow us to better understand the dynamics of outdoor access on the kinetics of cow gaits.

#### 3.2.3. Infrared Thermography

The results of the analyzed data regarding coronary band thermography revealed that the mean maximum temperature of lateral (LCB) and medial (MCB) claw declined from Pre-trial to Post-trial (P = 0.003), although the maximum temperature in lateral coronary band (LCB) was lower in Follow-up compared to Pre-trial (30.1 ± 0.29 °C, P = 0.001). This effect was not seen in the maximum temperature of the medial claw (i.e., MCB = 30.4 ± 0.25 °C, P = 0.16). Similar results were observed in the mean average temperature of the hoof coronary band (CB) and lateral and medial claws (LCB and MCB). The average temperature reduced 1.5 to 2 °C Pre-trial to Post-trial (CB P = 0.003, LCB P = 0.003 and MCB P = 0.0005) and Follow-up (CB P = 0.001, LCB P < 0.0001 and MCB P = 0.002). However, no effect of time was observed for the maximum temperature of the CB (P > 0.05; Table 7).

**Table 7.** Average ± standard errors of variables obtained from Infrared thermography from the dorsal view of the hoof (coronary band) followed by the statistical significance of treatments (EX1 and EX3) and time points (i.e., Pre-trial, Post-trial and Follow-up). Different letters in superscript indicate the significance.

| Variable<sup>1</sup> | Treatment Group |  | Time |  |  | P-value |  |  |
| --- | --- | --- | --- | --- | --- | --- | --- | --- |
|  | EX1 | EX3 | Pre-trial | Post-trial | Follow-up | TRT<sup>2</sup> | Time | TRT x Time |
| CB_Minimum Temperature (°C) | 21.1 ± 0.20 | 21.7 ± 0.20 | 22.1 ± 0.28<sup>a</sup> | 21.1 ± 0.27<sup>b</sup> | 21.0 ± 0.26<sup>b</sup> | 0.72 | 0.028 | 0.69 |
| CB_Maximum temperature (°C) | 31.6 ± 0.23 | 32.1 ± 0.23 | 32.5 ± 0.31 | 31.4 ± 0.29 | 31.8 ± 0.29 | 0.40 | 0.30 | 0.50 |
| CB_Maximum minus Minimum temperature (°C) | 10.6 ± 0.24 | 10.5 ± 0.24 | 10.4 ± 0.31 | 10.4 ± 0.29 | 10.9 ± 0.28 | 0.86 | 0.77 | 0.64 |
| CB_Average temperature (°C) | 27.9 ± 0.17 | 28.3 ± 0.17 | 29.1 ± 0.26<sup>a</sup> | 27.6 ± 0.25<sup>b</sup> | 27.7 ± 0.24<sup>b</sup> | 0.68 | 0.005 | 0.45 |
| CB_Standard deviation of temperature (°C) | 1.88 ± 0.05 | 1.88 ± 0.05 | 1.89 ± 0.05 | 1.78 ± 0.05 | 1.91 ± 0.05 | 0.85 | 0.75 | 0.61 |
| LCB_Minimum Temperature (°C) | 26.1 ± 0.19 | 26.4 ± 0.19 | 27.4 ± 0.32<sup>a</sup> | 26.1 ± 0.27<sup>b</sup> | 25.4 ± 0.25<sup>b</sup> | 0.78 | 0.001 | 0.41 |
| LCB_Maximum Temperature (°C) | 30.4 ± 0.22 | 30.8 ± 0.22 | 31.7 ± 0.29<sup>a</sup> | 30.0 ± 0.27<sup>b</sup> | 30.1 ± 0.26<sup>b</sup> | 0.81 | 0.001 | 0.26 |
| LCB_Maximum minus Minimum temperature (°C) | 4.35 ± 0.14 | 4.47 ± 0.15 | 4.42 ± 0.22<sup>ab</sup> | 3.84 ± 0.16<sup>a</sup> | 4.97 ± 0.19<sup>b</sup> | 0.35 | 0.002 | 0.25 |
| LCB_Average temperature (°C) | 29.0 ± 0.20 | 29.2 ± 0.20 | 30.3 ± 0.28<sup>a</sup> | 28.7 ± 0.26<sup>b</sup> | 28.4 ± 0.25<sup>b</sup> | 0.81 | 0.0002 | 0.29 |
| LCB_Standard deviation of temperature (°C) | 0.88 ± 0.03 | 0.87 ± 0.04 | 0.88 ± 0.05<sup>ab</sup> | 0.78 ± 0.03<sup>a</sup> | 0.97 ± 0.04<sup>b</sup> | 0.18 | 0.0008 | 0.08 |
| MCB_Minimum temperature (°C) | 25.7 ± 0.21 | 26.3 ± 0.22 | 27.1 ± 0.33<sup>a</sup> | 25.5 ± 0.29<sup>b</sup> | 25.4 ± 0.30<sup>b</sup> | 0.30 | 0.03 | 0.95 |
| MCB_Maximum temperature (°C) | 30.3 ± 0.21 | 30.8 ± 0.22 | 31.4 ± 0.35<sup>a</sup> | 29.9 ± 0.28<sup>b</sup> | 30.4 ± 0.25<sup>ab</sup> | 0.9 | 0.02 | 0.23 |
| MCB_Maximum minus minimum temperature (°C) | 4.59 ± 0.15 | 4.53 ± 0.15 | 4.28 ± 0.19 | 4.38 ± 0.18 | 5.03 ± 0.18 | 0.098 | 0.99 | 0.006 |
| MCB_Average temperature (°C) | 28.7 ± 0.21 | 29.1 ± 0.22 | 30.0 ± 0.34 <sup>a</sup> | 28.3 ± 0.27 <sup>b</sup> | 28.4 ± 0.25 <sup>b</sup> | 0.92 | 0.003 | 0.57 |
| MCB_Standard deviation of temperature (°C) | 0.94 ± 0.03 | 0.88 ± 0.35 | 0.88 ± 0.04 | 0.91 ± 0.04 | 0.94 ± 0.39 | 0.04 | 0.49 | 0.02 |

<sup>1</sup>Abbreviation: CB = coronary band temperature; LCB= lateral claw coronary band temperature; MCB= medial claw coronary band temperature;
<sup>2</sup>TRT = treatment

The sole temperature results showed an increasing trend of the maximum temperature of Z10 and Z0 from 26.0 ± 0.36 °C and 28.1 ± 0.41°C, respectively at Pre-trial to Post-trial (Z10: 28.2 ± 0.35 °C, P < 0.0001; Z0: 29.3 ± 0.40 °C, P = 0.07) and Follow-up (Z10: 29.0 ± 0.36 °C, P < 0.0001; Z0: 30.8 ± 0.41 °C, P < 0.0001). Similarly, the mean average temperatures of Z10 and Z0 increased from Pre-trial (Z10: 22.4 ± 0.33 °C; Z0: 23.8 ± 0.35 °C) to Post-trial (Z10: 25.2 ± 0.32 °C, P < 0.0001; Z0: 24.9 ± 0.34 °C, P = 0.02) and Follow-up (Z10: 25.6 ± 0.33 °C, P < 0.0001; Z0: 25.2 ± 0.35 °C, P = 0.002), with no change from Post-trial to Follow-up. Opposite to the results of the skin parts of the sole (Z10 and Z0), maximum and average temperature of the keratinized tissues (LZ4 and MZ4) showed 1.3 °C to 2.2 °C reduction from Pre-trial (maximum temperature = LZ4: 23.8 ± 0.30 °C, P= 0.0001; MZ4: 23.7 ± 0.32 °C, P = 0.005 and average temperature = LZ4: 22.1 ± 0.26 °C, P < 0.0001; MZ4: 22.8 ± 0.28 °C, P < 0.0001) to Follow-up. The maximum and average temperature of the LZ4 and MZ4 showed a non-significant increase from Pre-trial to Post-trial (P > 0.05) and a significant reduction from Post-trial to Follow-up (P < 0.05, Table 8). No effect of treatment or the interaction between time and treatment was observed for either maximum and average temperatures of the coronary band and sole views in all ROIs (Table S5 and Table S6 found in supplementary materials).

**Table 8.** Average ± standard errors of variables obtained from Infrared thermography form the plantar view of the hoof (sole) followed by the statistical significance of treatments (EX1 and EX3) and time points (i.e., Pre-trial, Post-trial and Follow-up). Different letters in superscript indicate the significance.

| Variable<sup>1</sup> | Treatment Group |  | Time |  |  | P-value |  |  |
| --- | --- | --- | --- | --- | --- | --- | --- | --- |
| | EX1 | EX3 | Pre-trial | Post-trial | Follow-up | TRT<sup>2</sup> | Time | TRT × Time |
| Z10_Maximum temperature | 27.8 ± 0.36 | 27.6 ± 0.36 | 26.0 ± 0.36<sup>a</sup> | 28.2 ± 0.35<sup>b</sup> | 29.0 ± 0.36<sup>b</sup> | 0.60 | 6.95e-10 | 0.12 |
| Z10_Minimum temperature | 19.3 ± 0.30 | 19.6 ± 0.29 | 17.7 ± 0.32<sup>a</sup> | 20.3 ± 0.31<sup>b</sup> | 20.4 ± 0.32<sup>b</sup> | 0.60 | 3.23e-08 | 0.12 |
| Z10_Average temperature | 24.4 ± 0.33 | 24.4 ± 0.32 | 22.4 ± 0.33<sup>a</sup> | 25.2 ± 0.32<sup>b</sup> | 25.6 ± 0.33<sup>b</sup> | 0.74 | 5.74e-10 | 0.13 |
| Z0_Maximum temp | 29.3 ± 0.43 | 29.4 ± 0.43 | 28.1 ± 0.41<sup>a</sup> | 29.3 ± 0.4<sup>a</sup> | 30.8 ± 0.41<sup>b</sup> | 0.58 | 7.69e-05 | 0.13 |
| Z0_Minimum temp | 17.4 ± 0.28 | 17.3 ± 0.27 | 17.3 ± 0.37<sup>a</sup> | 18.8 ± 0.31<sup>b</sup> | 16.0 ± 0.33<sup>c</sup> | 0.89 | 8.83e-07 | 0.62 |
| Z0_Average temp | 24.6 ± 0.35 | 24.7 ± 0.35 | 23.8 ± 0.35 | 24.9 ± 0.34 | 25.2 ± 0.35 | 0.85 | 0.017 | 0.17 |
| MZ4_Maximum temp | 23.4 ± 0.30 | 23.5 ± 0.30 | 23.7 ± 0.32<sup>b</sup> | 24.3 ± 0.31<sup>b</sup> | 22.4 ± 0.32<sup>a</sup> | 0.88 | 3.42e06 | 0.091 |
| MZ4_Minimum temp | 18.7 ± 0.23 | 18.7 ± 0.23 | 18.8 ± 0.25<sup>a</sup> | 19.8 ± 0.24<sup>b</sup> | 17.6 ± 0.25<sup>c</sup> | 0.96 | 1.1e09 | 0.22 |
| MZ4_Average temperature | 21.4 ± 0.27 | 21.5 ± 0.26 | 21.8 ± 0.28<sup>a</sup> | 22.5 ± 0.27<sup>a</sup> | 20.0 ± 0.28<sup>b</sup> | 0.75 | 1.79e-10 | 0.12 |
| LZ4_Maximum temp | 23.4 ± 0.28 | 23.2 ± 0.28 | 23.8 ± 0.30<sup>a</sup> | 24.2 ± 0.29<sup>a</sup> | 22.0 ± 0.30<sup>b</sup> | 0.98 | 2.27e-06 | 0.50 |
| LZ4_Minimum temp | 19.0 ± 0.21 | 18.9 ± 0.21 | 19.3 ± 0.23<sup>a</sup> | 20.0 ± 0.22<sup>a</sup> | 17.6 ± 0.23<sup>c</sup> | 0.69 | 2.41e-12 | 0.19 |
| LZ4_Average temperature | 21.5 ± 0.24 | 21.4 ± 0.24 | 22.1 ± 0.26<sup>a</sup> | 22.4 ± 0.25<sup>a</sup> | 19.9 ± 0.26<sup>b</sup> | 0.94 | 3.62e-11 | 0.30 |

<sup>1</sup>Abbreviation: Z10 = zone 10 temperature; Z0 = zone 0 temperature; MZ4 = zone 4 of medial claw temperature.
<sup>2</sup>TRT = treatment

Infrared thermography is a relatively novel technology in veterinary medicine, primarily used for detecting alterations in skin temperature due to pain and inflammation, before clinical signs appear (Alsaaod et al., 2015). Laminitis and claw lesions, in their early subclinical stages, can trigger immune responses, such as inflammation, which can be detected through an increase in coronary band temperature. (Nikkhah et al., 2005, Bobić et al., 2017, Arican et al., 2018). Our results depicted a reduction in coronary band temperature of the overall hoof and both medial and lateral claws for both treatment cows, which is in contrast to Nejati et al. (2024) which found no effect of outdoor access on the coronary band temperature. We believe that the discrepancy with our findings might be due to some differences in methodology. While in our study, the IRT images were taken in a room with controlled temperature to minimize the effect of ambient temperature (Alsaaod et al., 2014, Landgraf et al., 2014), the thermographs in Nejati et al. (2024) were taken in the barn, where the ambient temperature was not controlled and fluctuated through the data collection. In addition, we added the cow’s eye maximum, an indicator of the cow’s core body temperature, as a co-variable to adjust for the individual variation between cows.

Coronary band is a well-vascularized area of the hoof and its temperature can be affected by the circulation and the metabolic activity of the underlying tissue (Alsaaod et al., 2015). Some claw lesions, such as sole and white line hemorrhage, can be indicative of subclinical laminitis (Greenough and Vermunt, 1991, Stone, 2004, Passos et al., 2023). Laminitis is a painful condition in which inflammatory mediators such as histamine, interleukin-6, and lipopolysaccharides play a critical role in the pathogenesis of subclinical stages (Zhang et al., 2020, Passos et al., 2023) which might explain the correlation between the increase in locomotion score and elevation in coronary band temperature in cows (Rodríguez et al., 2016, Cramer et al., 2023). The reduction in coronary band temperature in our study might indicate that the provision of outdoor access as low as 1h/week could result in a reduction of inflammatory agents in blood circulation and subclinical laminitis and/or the pain associated with that. Further research on the serum concentration of these biomarkers before and after the provision of outdoor access and their relationship with coronary band temperature might be needed to corroborate our IRT results and better understand how outdoor access would affect subclinical laminitis.

The sole temperature in zone 4 (typical area for sole ulcers) exhibited a reduction at the Follow-up data collection point for both groups of treatment cows. Sole temperature has been shown to have a positive correlation with a cow’s gait score and a negative relationship with digital cushion thickness (Oikonomou et al., 2014, Rodríguez et al., 2016). Gard et al. (2015) reported an increase in digital cushion volume and surface after provision of exercise and more movement opportunities. Although the thickness of the digital cushion has not been investigated in our study, it can explain our results regarding zone 4 temperature, which may suggest that the provision of outdoor access might positively affect digital cushion development in cows, therefore inducing more comfortable walking and standing, as well as reducing the risk of lameness (Griffiths et al., 2020). However, another explanation would be that a reduction in zone 4 temperature might be due to an increase in the sole thickness. Although the sole thickness was not measured in our study, the increase in toe length and reduction of sole width (section <u>2.2.3</u>) might suggest the increase in the sole thickness of claws in the Follow-up data collection point.

Contrary to coronary band and zone 4, maximum and average temperature increased in zone 10 (i.e., the skin above claws) and zone 0 (the skin between claws) at both treatment groups. Provision of outdoor access has been shown to be related to increased risk of infectious diseases, including digital and interdigital dermatitis, which occurs in zone 10 and 0, respectively (Häggman and Juga, 2015, Moreira et al., 2019). Given that no infectious diseases were detected during this study, we hypothesize that walking on soil, grass, and natural substances in the exercise yard can cause minor injuries to the sole’s skin. On the other hand, Bielfeldt et al. (2005) suggested that exercise can positively affect claws by increasing the blood circulation in the area and transferring more nutrients and oxygen to live tissues. Both hypotheses could explain our observed increase in sole’s temperature in zones 0 and 10. Overall, our results on the hoof and sole thermography showed that increasing movement opportunity could have promising effects on the hoof health of cows; however, more research is needed to better understand the effect of outdoor access on blood circulation and anatomy of the hoof.

#### 3.2.4. Measuring Claw Conformation

Claw measurements showed that sole width, sole length, and claw length increased from Pre-trial (5.23 ± 0.04 cm, 8.92 ± 0.08 cm, and 8.03 ± 0.06 cm, respectively) to Post-trial (5.47 ± 0.04 cm, 9.46 ± 0.08 cm, and 8.43 ± 0.06 cm, P < 0.0001). At Follow-up, reductions in sole width (.29 ± 0.04 cm, P = 0.004) and sole length (9.35 ± 0.10 cm, P = 1.0) were observed while claw length increased from Post-trial to Follow-up (8.90 ± 0.06 cm, P <.0001). No effect of treatment or the interaction between time and treatment was observed for the mentioned variables. Similarly, there was no effect of time, treatment or their interaction for the claw angle measurements (Table 9 and Table S7 found in supplementary materials).

**Table 9.** Average ± standard errors of claw and hoof dimensions followed by the statistical significance of treatments (EX1 and EX3) and time points (i.e., Pre-trial, Post-trial and Follow-up). Different letters in superscript indicate the significance.

| Variable | Treatment Group |  | Time |  |  | P-value |  |  |
| --- | --- | --- | --- | --- | --- | --- | --- | --- |
| | EX1 | EX3 | Pre-trial | Post-trial | Follow-up | TRT<sup>1</sup> | Time | TRT × Time |
| Sole width (cm) | 5.35 ± 0.04 | 5.31 ± 0.04 | 5.23 ± 0.04<sup>a</sup> | 5.47 ± 0.04<sup>b</sup> | 5.29 ± 0.04<sup>a</sup> | 0.163 | 0.0003 | 0.23 |
| Sole length (cm) | 9.16 ± 0.08 | 9.33 ± 0.08 | 8.92 ± 0.08<sup>a</sup> | 9.46 ± 0.08<sup>b</sup> | 9.35 ± 0.10<sup>b</sup> | 0.68 | 0.006 | 0.78 |
| Claw length (cm) | 8.56 ± 0.07 | 8.35 ± 0.70 | 8.03 ± 0.06<sup>a</sup> | 8.43 ± 0.06<sup>b</sup> | 8.90 ± 0.06<sup>c</sup> | 0.028 | 3.33e-15 | 0.19 |
| Toe angle (cm) | 42.6 ± 0.56 | 42.2 ± 0.54 | 42.8 ± 0.61 | 42.3 ± 0.47 | 42.0 ± 0.36 | 0.45 | 0.19 | 0.47 |

<sup>1</sup>TRT = treatment

Our results showed an increase in claw length throughout of our study. This increase can be attributed to normal growth of claw horn tissue (Shearer and van Amstel, 2001). Our results are in line with Somers et al. (2005) who found positive effects of time on claw length regardless of the walking surface.

The increase in sole width at Post-trial may indicate more wear on the plantar surface when cows have more opportunity to move. These results support our results from 3D motion analysis and kinetic assessment that showed an increase in stance time, contact area and pressure applied by hooves after provision of outdoor access. However, at Follow-up, in which cows were not provided outdoor access for 8 weeks, sole width decreased. This reduction can be the result of lower opportunity for movement and less claw contact with the ground, which we hypothesize can lead to a higher growth rate of walls. It is important to note that this reduction is relatively small (about 2mm), as all the measurements were done after sliver trimming, which could also influence the reduction. Similar to claw length, sole length increase at Post-trial can be the result of normal growth of claw horn tissue (Shearer and van Amstel, 2001). We did not measure the growth and wear rate of claw horn tissue in this study. Although Somers et al. (2005) found that claw dimensions increase over time regardless of the flooring type, they did not find any impact of flooring type on the growth rate, wear rate, or shape of the claws. No difference in claw conformation was observed between our two treatment groups. Similar to our findings, Loberg et al. (2004) found no difference in the net growth of claws for cows with 1 and 2 days per week of outdoor access. However, they found the net growth in the diagonal (i.e., the distance from the tip of the toe to the proximal end of the heel) was smaller in cows with access to the outdoors (1, 2, and 7 days per week) compared to those without outdoor access. More studies on growth and wear rate, as well as additional measures of claw parameters, such as diagonal length or heel height, can provide better insight into how outdoor access and confinement would affect claw horn tissue.

Technologies have been applied to detect small changes in gait pattern and increase the accuracy of lesion detection at their early non-visible stages. Previous studies that compared visual locomotion scoring and some automated locomotion scoring systems showed that the sensitivity and specificity of these systems to recognize cows with lesions can be lower than human locomotion scoring (Bicalho et al., 2007, Anagnostopoulos et al., 2023). For example, Schlageter-Tello et al. (2018) compared a 3D automatic locomotion scoring system (3D ALS) and human locomotion scoring in the detection of cows with painful lesions. Their results showed that 3D automatic locomotion scoring had lower specificity and less success in detecting non-lame cows and had more false-positive. It should be considered that these technologies are still developing, and comparing them with clinical assessment methods, which are well developed to diagnose clinical signs, might cause low reliability. Although automated systems might not be prone to objectivity, tiredness, or distraction a human observer might have, the reason behind their poor performance in lameness detection might be due to focusing on limited gait characteristics (e.g., only back posture) or poor development of their algorithms (Schlageter-Tello et al., 2018, Logan et al., 2024). For example, when different kinematic traits of gait (i.e., walking speed, back arch, stride length, track up and head bob) were studied, it revealed that cows can mask signs of lameness when motivated, which might not be visible with the naked eye (Mokhtarnazif et al., 2020).

## 4. Conclusion

Although clinical assessments showed no effect of limited outdoor access on gait and hoof health of cows, the subclinical assessments revealed some positive effects. Cows with regular outdoor access, even as low as 1 and 3 hours per week, had longer stride and stance time and applied more pressure while standing suggesting they gain more locomotion comfort. Reduction of coronary band temperature could be interpreted as signs of reduction in pain and discomfort of subclinical conditions like laminitis and increase in sole and digital cushion thickness. Technologies revealed the subclinical effects; however, more studies will be needed to assess distribution of pressure within claws, hoof blood circulation and digital cushion thickness. This study faced some limitations including the need to adapt the technologies designed for humans (i.e., 3D motions analysis and pressure plates) for use in cattle, highlighting lack of knowledge on tailoring and designing these tools for bovine applications.

## 5. Note

This research was supported by NSERC, Novalait, Dairy Farmers of Canada, and Valacta through Vasseur’s Industrial Research Chair on the Sustainable Life of Dairy Cattle (IRCPJ/492639-2015), as well as a contribution from the Dairy Research Cluster 3 (Dairy Farmers of Canada and Agriculture and Agri-Food Canada) under the Canadian Agricultural Partnership AgriScience Program (Activity 11), as well as Vasseur’s NSERC Discovery Grant (#ID RGPIN-2019-04728). Equipment used in the validation study was funded through Vasseur’s Canada Foundation for Innovation’s John R. Evans Leaders Fund (#ID JELF-20258). Additional stipend funding was provided through Op+Lait, NSERC CREATE, and McGill Graduate Excellence Fellowship awards.

We thank Marjorie Cellier, Jasmine Muszik, Nazanin Sarmadi, Rachel van Vliet, Tania Wolfe and Véronique Boyer (Department of Animal Science, McGill University) for their assistance with data collection and manuscript editing. We would like to express our gratitude to Farzaneh Rahmdani (Department of computer science, Université du Québec à Montréal) for her help with statistical analysis. We also acknowledge the staff of the McGill University Macdonald Campus Dairy Complex (Sainte-Anne-de-Bellevue, QC, Canada) for their support in animal handling, farm management and animal husbandry.

