## Supplemental tables for "Discovering subclinical effects of limited outdoor access on gait and hoof health of cows housed in movement-restricted environments"

**Table S1.** Convergence of models for all the variables is shown here, the number 1 indicates the successful convergence and the number zero indicates that the model failed to converge. The astride (*) showed which model was chosen based on the Akaike Information Criteria (AIC) as the best-fitting model. (NA indicating that the model was not tested for the correlation structure)

| | Statistical model | | | |
|---|---|---|---|---|
| **Variable**[1] | No correlation structure | General correlation structure | Autoregressive of order 1 | Compound symmetry |
| **NRS** | 1 | 0* | 0 | 1 |
| **Swinging out** | 1* | 1 | 1 | 1 |
| **Back arch** | 1 | 0 | 0* | 1 |
| **Track-up** | 1 | 0 | 1* | 1 |
| **Joint Flexion** | 1* | 0 | 1 | 1 |
| **Asymmetric step** | 1* | 0 | 1 | 1 |
| **Reluctance to bear weight** | 0* | 0 | 1 | 0 |
| **Stride length** | 1 | 0* | 1 | 1 |
| **Stride time** | 1 | 0* | 1 | 1 |
| **Velocity** | 0 | 1 | 0* | 1 |
| **Stance time** | 1* | 1 | 1 | 1 |
| **Track-up X** | 1 | 0* | 1 | 1 |
| **Track-up XY** | 1* | 1 | 1 | 1 |
| **Hock angle (Min)** | 0 | 0 | 0* | 0 |
| **Hock angle (Max)** | 1 | 1 | 1* | 1 |
| **Hock angle range of motion (ROM)** | 0 | 0 | 0* | 0 |
| **Hock angle (Med)** | 1 | 0 | 0* | 0 |
| **Hock angle (Avr)** | 1 | 0 | 0* | 1 |

| | | | | |
|---|---|---|---|---|
| **Pressure Distribution** | 1* | 1 | 1 | 1 |
| **MPressure_SC** | 1 | 0 | 1* | 1 |
| **AVPressure_SC** | 1 | 0 | 1* | 1 |
| **MPressure_30Srec** | 1* | 0 | 0 | 1 |
| **MForce_30Srec** | 1 | 1 | 1* | 1 |
| **AVForce_30Srec** | 1* | 1 | 1 | 1 |
| **Contact_area_SC** | 1 | 1* | 1 | 1 |
| **Contact_area_30Srec** | 1* | 1 | 1 | 1 |
| **CB_Minimum Temperature** | 1 | 1 | 0* | 1 |
| **CB_Maximum minus Minimum temperature** | 1 | 0 | 0* | 0 |
| **CB_Maximum temperature** | 0 | 0* | 0 | 0 |
| **CB_Average temperature** | 0 | 0* | 0 | 0 |
| **CB_Standard deviation of temperature** | 1 | 1 | 0* | 1 |
| **LCB_Minimum Temperature** | 0 | 1* | 0 | 1 |
| **LCB_Maximum Temperature** | 0 | 0* | 0 | 1 |
| **LCB_Maximum minus Minimum temperature** | 1 | 1 | 1* | 1 |
| **LCB_Average temperature** | 0 | 0* | 0 | 0 |
| **LCB_Standard deviation of temperature** | 1 | 1 | 1* | 1 |
| **MCB_Minimum temperature** | 0 | 1 | 0 | 1* |
| **MCB_Maximum temperature** | 0 | 0 | 0 | 1* |
| **MCB_Maximum minus minimum temperature** | 1* | 1 | 1 | 1 |
| **MCB_Average temperature** | 0 | 0 | 0 | 1* |
| **MCB_Standard deviation of temperature** | 1 | 0 | 1 | 0* |

| | | | | |
|---|---|---|---|---|
| **Z10_Maximum temperature** | 1 | 1 | 1* | 1 |
| **Z10_Minimum temperature** | 0* | 1 | 0 | 1 |
| **Z10_Average temperature** | 1* | 1 | 1 | 1 |
| **Z0_Maximum temp** | 1 | 1 | 1* | 1 |
| **Z0_Minimum temp** | 1* | 0 | 0 | 1 |
| **Z0_Average temp** | 1* | 1 | 1 | 1 |
| **MZ4_Maximum temp** | 0* | 1 | 0 | 1 |
| **MZ4_Minimum temp** | 0* | 0 | 0 | 0 |
| **MZ4_Average temperature** | 1 | 0 | 1 | 0* |
| **LZ4_Maximum temp** | 1* | 1 | 1 | 1 |
| **LZ4_Minimum temp** | 0* | 0 | 0 | 0 |
| **LZ4_Average temperature** | 0* | 0 | 1 | 0 |
| **Sole width** | 0* | 0 | 1 | 0 |
| **Sole length** | 1* | 1 | 1 | 1 |
| **Claw length** | 1 | 0 | 0* | 1 |
| **Toe angle** | 1* | 0 | 1 | 0 |
| **Lesion score** | 1* | NA | NA | NA |
| **Lesion prevalence** | 1* | NA | NA | NA |

**[1]Abbreviation: Min = minimum, Max = Maximum, Med= Median Avr = average, SC= Screenshot, 30Srec= 30 seconds recording, MPressure = maximum pressure, AVPressure = Average pressure, MForce= Maximum force, AVForce = average force, CB = coronary band temperature, LCB= lateral claw coronary band temperature, MCB= medial claw coronary band temperature, Z10 = zone 10 temperature, Z0 = zone 0 temperature, MZ4 = zone 4 of medial claw temperature**

**Table S2.** Average ± standard errors of overall gait (NRS) and gait attributes scores based on the interaction between the two treatment groups (i.e., EX1 and EX3) and each data collection point (i.e., Pre-trial, Post-trial and Follow-up). The P-value shows the interaction effect.

| Variable | Treatment | Pre-trial | Post-trial | Follow-up | P-value |
|---|---|---|---|---|---|
| **NRS** | EX1 | 2.16 ± 0.16 | 1.84 ± 0.16 | 1.91 ± 0.16 | 0.97 |
| | EX3 | 2.02 ± 0.16 | 1.75 ± 0.16 | 1.75 ± 0.16 | |
| **Swing out** | EX1 | 1.25 ± 0.20 | 1.16 ± 0.20 | 1.09 ± 0.21 | 0.29 |
| | EX3 | 1.48 ± 0.20 | 0.98 ± 0.20 | 1.39 ± 0.20 | |
| **Back arch** | EX1 | 0.55 ± 0.21 | 0.63 ± 0.21 | 0.89 ± 0.22 | 0.45 |
| | EX3 | 0.71 ± 0.21 | 0.44 ± 0.21 | 0.62 ± 0.21 | |
| **Tracking up** | EX1 | 0.96 ± 0.22 | 0.62± 0.21 | 0.83 ± 0.22 | 0.11 |
| | EX3 | 0.93 ± 0.22 | 0.84 ± 0.21 | 0.57 ± 0.21 | |
| **Joint flexion** | EX1 | 1.76 ± 0.16 | 1.43 ± 0.18 | 1.33 ± 0.25 | 0.19 |
| | EX3 | 1.35 ± 0.16 | 1.62 ± 0.18 | 1.27 ± 0.25 | |
| **Asymmetric step** | EX1 | 0.89 ± 0.19 | 0.89 ± 0.19 | 0.99 ± 0.21 | 0.83 |
| | EX3 | 0.87 ± 0.19 | 0.73 ± 0.19 | 0.73 ± 0.20 | |
| **Reluctance to bear weight** | EX1 | 0.78 ± 0.22 | 0.69 ± 0.22 | 0.72 ± 0.23 | 0.77 |
| | EX3 | 0.87 ± 0.22 | 0.55 ± 0.22 | 0.50 ± 0.22 | |

**Table S3.** Average ± standard errors of overall gait score (NRS) and gait attributes followed by the statistical significance of treatments (EX1 and EX3) and time points (i.e., Pre-trial, Post-trial and Follow-up).

| Variable | Treatment Group | | Time | | | P-value | | |
|---|---|---|---|---|---|---|---|---|
| | EX1 | EX3 | Pre-trial | Post-trial | Follow-up | Treatment | Time | Treatment × Time |
| NRS | 1.97 ± 0.10 | 1.84 ± 0.10 | 2.09 ± 0.11 | 1.79 ± 0.11 | 1.83 ± 0.11 | 0.55 | 0.26 | 0.97 |
| Swing out | 1.17 ± 0.15 | 1.29 ± 0.15 | 1.37 ± 0.14 | 1.07 ± 1.14 | 1.24 ± 0.14 | 0.42 | 0.80 | 0.29 |
| Back arch | 0.69 ± 0.15 | 0.59 ± 0.15 | 0.63 ± 0.15 | 0.54 ± 0.15 | 0.75 ± 0.15 | 0.58 | 0.45 | 0.45 |
| Tracking up | 0.80 ± 0.18 | 0.78 ± 0.18 | 0.94 ± 0.15 | 0.73 ± 0.15 | 0.70 ± 0.15 | 0.93 | 0.20 | 0.11 |
| Joint flexion | 1.51 ± 0.13 | 1.41 ± 0.13 | 1.55 ± 0.11 | 1.53 ± 0.13 | 1.30 ± 0.18 | 0.07 | 0.27 | 0.19 |
| Asymmetric step | 0.93 ± 0.12 | 0.78 ± 0.12 | 0.88 ± 0.14 | 0.81 ± 0.14 | 0.86 ± 0.14 | 0.93 | 0.91 | 0.83 |
| Reluctance to bear weight | 0.73 ± 0.13 | 0.64 ± 0.13 | 0.82 ± 0.15 | 0.62 ± 0.15 | 0.61 ± 0.16 | 0.78 | 0.95 | 0.77 |

**Table S4.** Average ± standard errors of variables obtained from 3D motion analysis of gait based on the interaction between the two treatment groups (i.e., EX1 and EX3) and each data collection point (i.e., Pre-trial, Post-trial and Follow-up). The P-value shows the interaction effect.

| Variable[1] | Treatment | Pre-trial | Post-trial | Follow-up | P-value |
|---|---|---|---|---|---|
| **Stride length (cm)** | EX1 | 161 ± 1.37 | 164 ± 1.40 | 162 ± 1.42 | 0.54 |
| | EX3 | 159 ± 1.30 | 160 ± 1.30 | 160 ± 1.30 | |
| **Stride time (s)** | EX1 | 1.07 ± 0.02 | 1.14 ± 0.02 | 1.09 ± 0.02 | 0.37 |
| | EX3 | 1.10 ± 0.02 | 1.12 ± 0.02 | 1.09 ± 0.02 | |
| **Velocity (cm/s)** | EX1 | 152 ± 2.29 | 145 ± 2.38 | 150 ± 2.41 | 0.36 |
| | EX3 | 145 ± 2.17 | 144 ± 2.17 | 147 ± 2.17 | |
| **Stance time (cm)** | EX1 | 0.69 ± 0.01 | 0.74 ± 0.01 | 0.70 ± 0.01 | 0.17 |
| | EX3 | 0.71 ± 0.01 | 0.73 ± 0.01 | 0.71 ± 0.01 | |
| **Track-up X (cm)** | EX1 | -1.29 ± 1.56 | 0.70 ± 1.58 | -0.82 ± 1.58 | 0.02 |
| | EX3 | 0.01 ± 1.47 | 0.23 ± 1.47 | 2.05 ± 1.47 | |
| **Track-up XY (cm)** | EX1 | 10.02 ± 0.83 | 10.61 ± 0.83 | 8.92 ± 0.92 | 0.08 |
| | EX3 | 10.03 ± 0.79 | 8.39 ± 0.78 | 9.87 ± 0.83 | |
| **Hock angle (Min)** | EX1 | 128 ± 1.08 | 126 ± 1.12 | 128 ± 1.12 | 0.48 |
| | EX3 | 128 ± 1.02 | 128 ± 1.02 | 130 ± 1.02 | |
| **Hock angle (Max)** | EX1 | 163 ± 1.04 | 162 ± 1.07 | 165 ± 1.19 | 0.48 |
| | EX3 | 162 ± 0.98 | 162 ± 0.98 | 164 ± 1.09 | |
| **Hock angle range of motion (ROM)** | EX1 | 35.8 ± 0.91 | 35.9 ± 0.95 | 36.6 ± 0.96 | 0.30 |
| | EX3 | 36.1 ± 0.86 | 33.9 ± 0.86 | 34.6 ± 0.86 | |
| **Hock angle (Med)** | EX1 | 147 ± 1.07 | 145 ± 1.10 | 147 ± 1.10 | 0.69 |
| | EX3 | 147 ± 1.01 | 146 ± 1.01 | 147 ± 1.01 | |
| **Hock angle (Avr)** | EX1 | 149 ± 1.07 | 148 ± 1.11 | 150 ± 1.12 | 0.16 |
| | EX3 | 150 ± 1.02 | 148 ± 1.02 | 148 ± 1.02 | |

[1]**Abbreviation:**

**Min = minimum, Max = Maximum, Med = Median Avr = average**

**Table S5.** Average ± standard errors of variables obtained from Infrared thermography from the dorsal view of the hoof (coronary band) based on the interaction between the two treatment groups (i.e., EX1 and EX3) and each data collection point (i.e., Pre-trial, Post-trial and Follow-up). The P-value shows the interaction effect. Different letters in superscript indicate the significance.

| Variable[1] | Treatment | Pre-trial | Post-trial | Follow-up | P-value |
|---|---|---|---|---|---|
| CB_Minimum Temperature | EX1 | 22.0 ± 0.39 | 20.7 ± 0.39 | 20.6 ± 0.37 | 0.69 |
| | EX3 | 22.2 ± 0.40 | 21.4 ± 0.37 | 21.4 ± 0.36 | |
| CB_Maximum temperature | EX1 | 32.2 ± 0.43 | 31.4 ± 0.42 | 31.2 ± 0.40 | 0.50 |
| | EX3 | 32.7 ± 0.44 | 31.6 ± 0.40 | 32.1 ± 0.40 | |
| CB_Maximum minus Minimum temperature | EX1 | 10.4 ± 0.43 | 10.6 ± 0.42 | 10.8 ± 0.40 | 0.64 |
| | EX3 | 10.5 ± 0.44 | 10.1 ± 0.40 | 11.0 ± 0.40 | |
| CB_Average temperature | EX1 | 28.9 ± 0.36 | 27.6 ± 0.35 | 27.1 ± 0.34 | 0.45 |
| | EX3 | 29.1 ± 0.37 | 27.8 ± 0.34 | 28.1 ± 0.33 | |
| CB_Standard deviation of temperature | EX1 | 1.89 ± 0.08 | 1.84 ± 0.07 | 1.90 ± 0.06 | 0.61 |
| | EX3 | 1.87 ± 0.08 | 1.74 ± 0.07 | 1.92 ± 0.06 | |
| LCB_Minimum temperature | EX1 | 27.3 ± 0.43 | 26.2 ± 0.49 | 25.0 ± 0.36 | 0.41 |
| | EX3 | 27.5 ± 0.45 | 25.9 ± 0.37 | 25.8 ± 0.35 | |
| LCB_Maximum Temperature | EX1 | 31.8 ± 0.40 | 29.9 ± 0.39 | 29.6 ± 0.38 | 0.26 |
| | EX3 | 31.6 ± 0.42 | 30.0 ± 0.38 | 30.6 ± 0.37 | |
| LCB_Maximum minus Minimum temperature | EX1 | 4.62 ± 0.30 | 3.59 ± 0.24 | 4.84 ± 0.28 | 0.25 |
| | EX3 | 4.22 ± 0.31 | 4.08 ± 0.22 | 5.11 ± 0.27 | |
| LCB_Average temperature | EX1 | 30.4 ± 0.38 | 28.7 ± 0.37 | 28.0 ± 0.36 | 0.29 |
| | EX3 | 30.2 ± 0.39 | 28.6 ± 0.36 | 28.7 ± 0.35 | |
| LCB_Standard deviation of temperature | EX1 | 0.94 ± 0.07 | 0.73 ± 0.05 | 0.97 ± 0.06 | 0.08 |
| | EX3 | 0.82 ± 0.07 | 0.83 ± 0.05 | 0.97 ± 0.06 | |
| MCB_Minimum temperature | EX1 | 26.7 ± 0.46 | 25.2 ± 0.42 | 25.1 ± 0.44 | 0.95 |
| | EX3 | 27.4 ± 0.47 | 25.9 ± 0.39 | 25.6 ± 0.43 | |
| MCB_Maximum temperature | EX1 | 31.4 ± 0.49 | 29.8 ± 0.40 | 29.7 ± 0.37 | 0.23 |
| | EX3 | 31.3 ± 0.50 | 30.0 ± 0.38 | 31.1 ± 0.35 | |
| MCB_Maximum minus minimum temperature | EX1 | 4.59 ± 0.26 | 4.59 ± 0.27 | 4.60 ± 0.26 | 0.006 |
| | EX3 | 3.96 ± 0.27[a] | 4.17 ± 0.25[a] | 5.45 ± 0.25[b] | |
| MCB_Average temperature | EX1 | 30.0 ± 0.47 | 28.2 ± 0.39 | 27.9 ± 0.36 | 0.57 |
| | EX3 | 30.1 ± 0.49 | 28.5 ± 0.37 | 28.8 ± 0.35 | |
| MCB_Standard deviation of temperature | EX1 | 0.97 ± 0.06 | 0.97 ± 0.06 | 0.89 ± 0.06 | 0.02 |
| | EX3 | 0.80 ± 0.06 [a] | 0.85 ± 0.06[ab] | 0.99 ± 0.88 [b] | |

**[1]Abbreviation: CB = coronary band temperature, LCB= lateral claw coronary band temperature, MCB= medial claw coronary band temperature**

**Table S6.** Average ± standard errors of variables obtained from Infrared thermography from the plantar view of the hoof (sole) based on the interaction between the two treatment groups (i.e., EX1 and EX3) and each data collection point (i.e., Pre-trial, Post-trial and Follow-up). The P-value shows the interaction effect.

| Variable[1] | Treatment | Pre-trial | Post-trial | Follow-up | P-value |
|---|---|---|---|---|---|
| **Z10_Maximum temperature** | EX1 | 25.8 ± 0.48 | 28.8 ± 0.48 | 28.9 ± 0.48 | 0.12 |
| | EX3 | 26.2 ± 0.50 | 27.6 ± 0.48 | 27.6 ± 0.48 | |
| **Z10_Minimum temperature** | EX1 | 17.5 ± 0.42 | 20.6 ± 0.42 | 19.9 ± 0.45 | 0.12 |
| | EX3 | 17.8 ± 0.44 | 20.1 ± 0.42 | 21.0 ± 0.44 | |
| **Z10_Average temperature** | EX1 | 22.3 ± 0.44 | 25.5 ± 0.44 | 25.2 ± 0.47 | 0.13 |
| | EX3 | 22.5 ± 0.47 | 24.8 ± 0.44 | 25.9 ± 0.46 | |
| **Z0_Maximum temp** | EX1 | 27.9 ± 0.56 | 29.8 ± 0.55 | 30.4 ±0.58 | 0.13 |
| | EX3 | 28.3 ± 0.58 | 28.8 ± 0.55 | 31.2 ±0.57 | |
| **Z0_Minimum temp** | EX1 | 17.2 ± 0.49 | 19.1 ± 0.46 | 15.9 ± 0.47 | 0.62 |
| | EX3 | 17.3 ± 0.51 | 18.5 ± 0.43 | 16.1 ± 0.45 | |
| **Z0_Average temp** | EX1 | 23.7 ± 0.47 | 25.2 ± 0.50 | 24.8 ± 0.50 | 0.17 |
| | EX3 | 23.8 ± 0.50 | 24.6 ± 0.47 | 25.6 ± 0.50 | |
| **MZ4_Maximum temp** | EX1 | 23.7 ± 0.43 | 24.7 ± 0.43 | 22.0 ± 0.45 | 0.091 |
| | EX3 | 23.8 ± 0.45 | 23.9 ± 0.43 | 22.9 ± 0.44 | |
| **MZ4_Minimum temp** | EX1 | 18.9 ± 0.33 | 20.0 ± 0.33 | 17.3 ± 0.35 | 0.22 |
| | EX3 | 18.8 ± 0.35 | 19.5 ± 0.33 | 17.9 ± 0.34 | |
| **MZ4_Average temperature** | EX1 | 21.8 ± 0.37 | 22.8 ± 0.37 | 19.6 ± 0.39 | 0.12 |
| | EX3 | 21.9 ± 0.39 | 22.2 ± 0.37 | 20.4 ± 0.39 | |
| **LZ4_Maximum temp** | EX1 | 23.8 ± 0.40 | 24.5 ± 0.40 | 22.0 ± 0.42 | 0.50 |
| | EX3 | 23.8 ± 0.42 | 23.9 ± 0.40 | 22.1 ± 0.42 | |
| **LZ4_Minimum temp** | EX1 | 19.3 ± 0.31 | 20.3 ± 0.31 | 17.4 ± 0.33 | 0.19 |
| | EX3 | 19.2 ± 0.32 | 19.7 ± 0.31 | 17.8 ± 0.32 | |
| **LZ4_Average temperature** | EX1 | 22.1 ± 0.35 | 22.7 ± 0.35 | 19.7 ± 0.37 | 0.30 |
| | EX3 | 22.1 ± 0.37 | 22.1 ± 0.35 | 20.0 ± 0.36 | |

**[1]Abbreviation:**

**Z10 = zone 10 temperature, Z0 = zone 0 temperature, MZ4 = zone 4 of medial claw temperature,**

**Table S7.** Average ± standard errors of claw and hoof dimensions based on the interaction between the two treatment groups (i.e., EX1 and EX3) and each data collection point (i.e., Pre-trial, Post-trial and Follow-up). The P-value shows the interaction effect.

| Variable | Treatment | Pre-trial | Post-trial | Follow-up | P-value |
|---|---|---|---|---|---|
| **Sole width** | EX1 | 5.20 ± 0.06 | 5.50 ± 0.60 | 5.36 ± 0.07 | 0.23 |
| | EX3 | 5.26 ± 0.60 | 5.44 ± 0.62 | 5.23 ± 0.06 | |
| **Sole length** | EX1 | 8.88 ± 0.12 | 9.37 ± 0.11 | 9.23 ± 0.14 | 0.78 |
| | EX3 | 8.95 ± 0.12 | 9.55 ± 0.11 | 9.48 ± 0.13 | |
| **Claw length** | EX1 | 8.17 ± 0.09 | 8.56 ± 0.09 | 8.94 ± 0.09 | 0.19 |
| | EX3 | 7.89 ± 0.09 | 8.31 ± 0.08 | 8.86 ± 0.08 | |
| **Toe angle** | EX1 | 43.3 ± 0.87 | 42.7 ± 0.69 | 41.8 ± 0.54 | 0.47 |
| | EX3 | 42.3 ± 0.88 | 42.0 ± 0.65 | 42.2 ± 0.51 | |
